# Fatal consequences of feline coronavirus infection are associated with virus persistence and a distinct adaptive immune repertoire

**DOI:** 10.1101/2025.10.02.679983

**Authors:** Thomas K. Hiron, Sara Falcone, The MASCOT Consortium, Anja Kipar, Melanie J. Hezzell, Chris A. O’Callaghan, Emi N. Barker, Lucy J. Davison

## Abstract

The pathogenesis of feline infectious peritonitis (FIP), arising in a minority of cats infected with feline coronavirus (FCoV), is complex and incompletely understood. Without extended use of direct-acting antivirals FIP is invariably fatal, but there is potential for the emergence of anti-viral resistance. To understand host and viral factors associated with FIP, multiple tissues from cats with and without FIP were subjected to RNA sequencing (RNA-seq), and targeted sequencing of the T cell receptor (TCR) repertoire was conducted for mesenteric lymph nodes in a larger cohort. Samples from cats with FIP demonstrated higher expression of genes involved in type I interferon and proinflammatory cytokine signalling, as well as the adaptive immune response, and expression of these genes was highly correlated with FCoV abundance. Analysis of FCoV genomic variation across tissues revealed dynamic within-host evolution of FCoV, and identified distinct mutations associated with systemic virus spread both within and among cats. Assembly of TCR and B-cell receptor (BCR) sequences identified changes in the immune repertoire associated with FIP, highlighting the polyclonality and ineffectiveness of the immune response to FCoV in cats with FIP, and revealing the presence of potentially protective TCR clonotypes in cats without FIP. Together, these results represent the first analysis of immune repertoire in any feline infectious disease and demonstrate naturally occurring within-host evolution of FCoV. These novel insights into a life-threatening systemic coronaviral disease have the potential to transform therapeutic approaches in FIP.

## Introduction

Feline Infectious Peritonitis (FIP) is an uncommon manifestation of infection with feline coronavirus (FCoV), characterised by immune dysregulation, hyperglobulinaemia, and granulomatous inflammatory processes including vasculitis (1). FCoV is a positive-sense single-stranded RNA virus belonging to the species *Alphacoronavirus suis*. Wild and domestic felids are susceptible to infection, typically resulting in mild self-limiting diarrhoea, but in some cats FIP develops from a delayed and dysregulated systemic response to infection (2) and is fatal without timely direct-acting anti-viral treatment (3–5). However, the cost of treatment is significant, not all cats respond, and reports have emerged of drug-associated side-effects and disease relapse in treated cats (6,7). It is also uncertain whether the selective pressure of anti-viral therapy will drive the emergence of drug resistance. The *intra vitam* diagnosis of FIP can be complicated due to the wide spectrum of clinical signs, ranging from ‘dry’ granulomatous disease to ‘wet’ effusive disease, and a comprehensive set of diagnostic guidelines has been developed by the European Advisory Board on Cat Diseases (ABCD) (8).

It remains unclear why only a minority (5-12%) of FCoV-infected cats develop FIP, although purebred cats and those in multi-cat households appear disproportionately affected (9,10). An examination of host-related factors in Birman cats by genome-wide association study (GWAS) revealed FIP-associated variants in and near genes involved in immune regulation and apoptosis, but no variants were fully concordant with the disease phenotype (11). Additional efforts to characterise the role of the host immune response in FIP have included RNA-seq of mesenteric lymph nodes (MLN) from cats of varying FCoV infection status, highlighting the involvement of the type I interferon response in FIP (12). Coronavirus-mediated FIP-like syndromes have been observed in other species, in ferrets (i.e. ferret systemic coronavirus disease (13)), and in dogs infected with the CB/05 strain of canine coronavirus (CCoV) (14), and similarities have been noted between FIP and certain aspects of COVID-19, caused by SARS-CoV-2, in humans (15).

Hypergammaglobulinaemia is commonly present in FIP-affected cats, and the host adaptive immune response during FCoV infection appears to influence progression to FIP (16,17). Experimentally, immunosuppressed cats demonstrate increased susceptibility to FIP, whilst some cats are resistant to experimental infection with known pathogenic FCoV strains (18). Broadly, cell-mediated (T cell) immunity is considered to be protective, and humoral (B cell) immunity is considered to be detrimental, and antibody-dependent enhancement has been proposed as one mechanism for the historic inefficacy of vaccines designed to protect against FIP (17,19,20). The antigen-specificity of the aberrant adaptive immune response in FIP has not been determined.

In recent years, protocols have been developed for targeted sequencing of T-cell receptor (TCR) and B-cell receptor (BCR) sequences comprising an individual’s immune repertoire, and changes in the immune repertoire have been associated with a range of human immune-mediated diseases (including COVID-19) (21,22). Such approaches have yet to be applied to feline disease, as the full extent of the feline TCR genomic locus has only recently been described (23), and the BCR heavy chain locus remains unresolved in the most commonly used feline reference genome (felCat9.0).

In addition to host factors, viral factors also play an important role in the development of FIP. Numerous studies have identified mutations in FCoV genes suspected to alter tropism from enterocytes to monocytes/macrophages, enabling the virus to spread to multiple tissues from the blood and to induce granulomatous inflammatory processes (24). However no single critical mutation has been identified, and diverse FCoV genomic sequences within FCoV-infected cats have been reported (25). Two FCoV biotypes are described: feline enteric coronavirus (FECV) and feline infectious peritonitis virus (FIPV), with the ‘internal mutation’ hypothesis proposing that *de novo* mutations in low virulence FECV genomes within an infected host give rise to FIP (26). An alternative hypothesis, known as ‘circulating virulent-avirulent FCoV’, posits that pathogenic and non-pathogenic strains of FCoV are in constant circulation within feline populations and development of FIP results from horizontal transmission of pathogenic strains (27). The internal mutation hypothesis best explains the majority of isolated cases of FIP, while the ‘circulating virulent-avirulent FCoV’ hypothesis best explains the infrequent outbreaks of FIP, including the recent high mortality outbreak in Cyprus (28).

Two serotypes of FCoV have been identified from cases of spontaneous infection, referred to as FCoV-1 and FCoV-2. Phylogenetic analysis indicates that FCoV-2 has arisen on multiple occasions as a consequence of recombination events between canine coronavirus (CCoV), also belonging to *Alphacoronavirus suis*, and FCoV-1 (29). This recombination event includes receipt of the CCoV S gene, encoding the spike protein required for viral entry. A FCoV-2 strain is associated with the Cypriot FIP outbreak, FCoV-23, in which the S protein shows 97% sequence identity to CCoV CB/05 (28). Coronaviruses belonging to *Alphacoronavirus suis* infect multiple species, and the potential for recombination between these closely related viruses has caused significant concern, with recent evidence of zoonotic transmission of recombinant strains (30,31). Currently, the majority of clinical FIP cases are associated with FCoV-1 (32–35), but most *in vitro* studies of FCoV to date have been conducted using FCoV-2 strains, due to the relative ease of culture compared with FCoV-1 (24).

Within the various quasispecies identified in FCoV-infected cats, specific mutations have been associated with FIPV, such as the M1058L and S1060A substitutions in a putative fusion domain of the FCoV spike protein (36). However, subsequent studies identified FCoV isolates with these mutations in cats with systemic infection, but without FIP, leading to the hypothesis that whilst such mutations may reflect a change in tropism or capacity to replicate in monocytes, in isolation they are not sufficient for development of FIP (32). Additional mutations in the furin cleavage site at the S1/S2 domain boundary of the FCoV spike protein can also distinguish FIPV from FECV in some cats (37). The polybasic S-R-R-S/A-R-R-S motif is highly conserved in FECV, while at least one residue in this motif is mutated generally in FIPV, suggesting that altered cleavage by host cell proteases (e.g. furin) may contribute to FIP pathogenesis. The S1/S2 cleavage site is notably absent from the S gene in FCoV-2. Targeted sequencing of FCoV accessory genes, encoded by ORF3abc and ORF7ab, has also identified candidate FIP-associated mutations, but the precise function of the encoded accessory proteins is largely unknown and the mutations are not reliably associated with disease (24).

Here, we report the use of RNA-sequencing to characterise both viral sequences and host immune responses in multiple tissues by the Mapping Animal Susceptibility to Coronavirus: Outcomes and Transcriptomics (MASCOT) consortium. We identified differentially expressed genes in liver, lung and MLN from FIP and non-FIP cats, and assembled full-length FCoV genomes from different tissues from multiple cats with quantitative analysis of intrahost FCoV genetic variation. Finally, we performed the first analysis of the B-cell and T-cell immune repertoire of cats with and without FIP, including clonotypes exclusive to either FIP or non-FIP cats. This study is the first to investigate the host transcriptomic response to FIP in multiple tissues in an unbiased manner, the first to explore the immune repertoire in any feline infectious disease, and the first to characterise and analyse intra-host variation of FCoV by RNA-seq. Our results represent the most comprehensive analysis to date of viral and host immune factors in FIP and will be of particular relevance in the design of novel preventative and therapeutic strategies in this potentially fatal disease.

## Results

### Multi-tissue RNA-seq of cats with or without FIP

Feline tissue samples were selected from the Bristol FIP Biobank (32) based on clinical information, histopathological analysis of tissues, and presence of FCoV antigen within intralesional macrophages confirmed by immunohistochemistry in at least one tissue (Fig. S1a). Samples from liver, lung, and mesenteric lymph node (MLN) were prioritised, as these tissues have been found to contain pyogranulomatous lesions in cats with FIP (38,39). Two cats with a diagnosis of a focal renal form of FIP, of which one had concurrent neurological FIP, were included for comparison. Control cats were selected based on availability of archived tissues from cats with a definitive alternative diagnosis, and IHC and histopathological changes not consistent with FIP. RNA was extracted from all selected FIP and control samples, and resulting total RNA was used for RNA-seq (Fig. 1a). The sample numbers for each group (FIP and control), and each tissue (liver, lung, and MLN) included in the RNA-seq are shown in Fig. 1b, and a full list of samples, including clinical and histopathological observations, RT-qPCR, and IHC results, is shown in Supplementary Table S1. All tissue RNA samples were sequenced as Illumina paired-end reads to a mean depth of 161,221,417. For all samples, >93% reads were mapped to the feline reference genome (Felis_catus_9.0/felCat9), and >81% were uniquely mapped to a single locus (Fig. S1b). Alignments were filtered to retain only uniquely mapped, properly paired reads, and gene-level counts were obtained. Principal component analysis (PCA) of normalised counts reflected distinct transcriptomes for each tissue (Fig. 1c), and PCA of samples from each tissue separately indicated alterations of the transcriptome in FIP samples compared to control samples (Fig. S1c-e). Differentially expressed genes (DEGs) between FIP and control samples (adjusted p-value < 0.05) were detected in all tissues, with 1,430 in the liver, 118 in the lung and 826 in the MLN (Fig. 1d). Lists of DEGs from each tissue are available in Supplementary Tables S2-S4.

**Figure 1.**
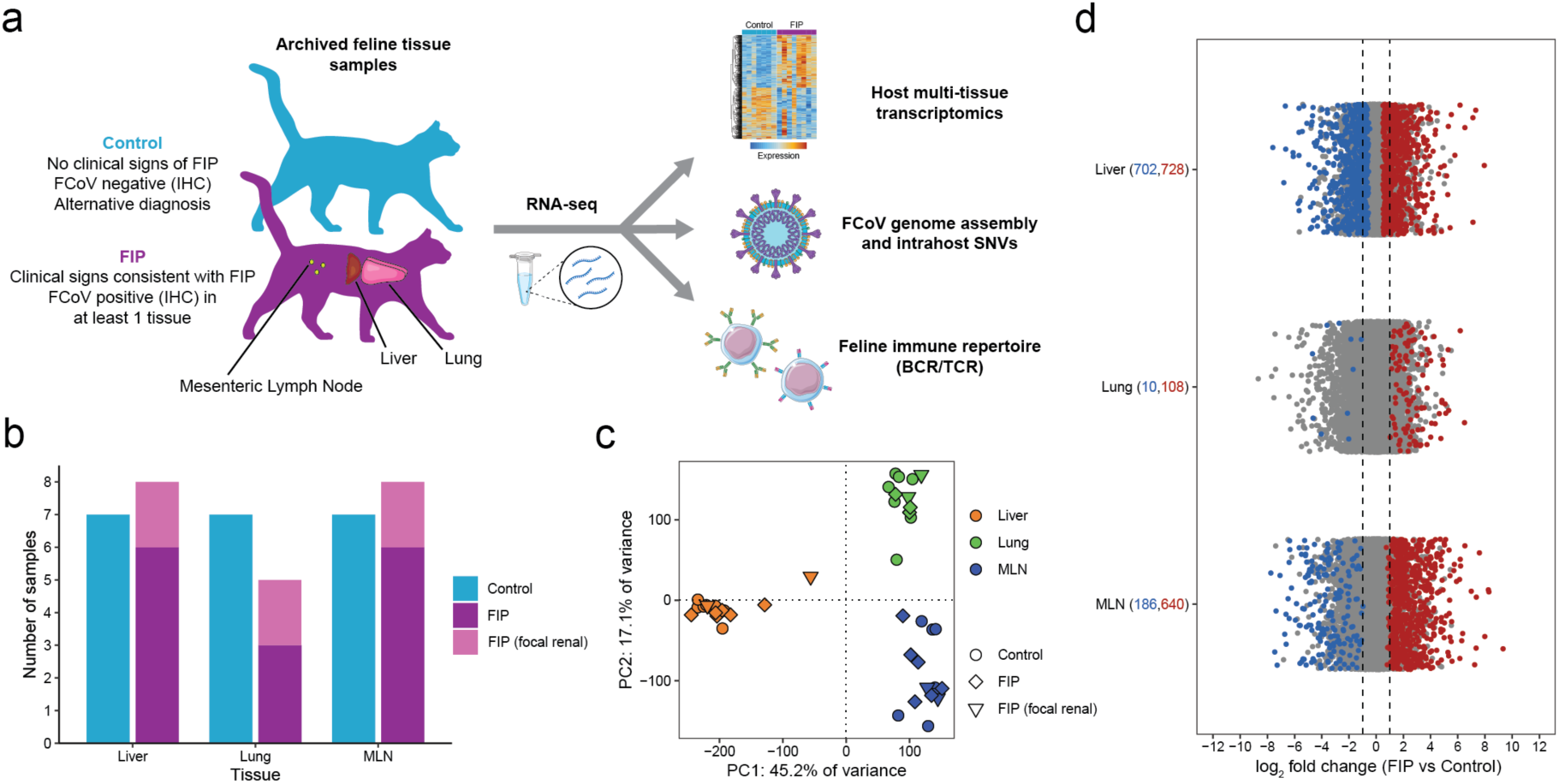
Multi-tissue RNA-seq of FIP cases and controls. (a) Overview of the study, with details of sample selection and analyses conducted using RNA-seq samples. FCoV – feline coronavirus, IHC – immunohistochemistry, SNV – single nucleotide variant, BCR – B-cell receptor, TCR – T-cell receptor. (b) Number of FIP and Control samples analysed for each tissue. FIP samples are further divided into those with more generalised FIP clinical signs, and those with a focal renal form of FIP. MLN – mesenteric lymph node. (c) Results of principal component analysis (PCA) of all tissue samples subjected to RNA-seq as part of the study. Each point represents an individual sample. The shape of each point corresponds to the sample group, and the colour of each point corresponds to the tissue from which the sample was taken. (d) Results of differential gene expression analysis showing log2 fold changes between FIP and Control samples for all tissues. Grey points represent genes with non-significant changes in expression (adjusted p-value > 0.05), and red and blue points represent significantly (adjusted p-value < 0.05) up- and down-regulated genes in FIP samples, respectively. Dashed black lines mark a two-fold increase or decrease in expression in FIP samples compared to Control samples. The number of differentially expressed genes with increased or decreased expression in FIP samples are shown in brackets in red and blue, respectively.

### Activation of type I IFN response, inflammatory cytokines and adaptive immunity in FIP

Comparison of the DEGs identified in each tissue revealed a subset of 198 genes with higher expression in multiple tissues from cats with FIP, of which 53 genes were more highly expressed in all 3 tissues analysed (Fig. 2a). Genes with higher expression in FIP common to all tissues included those with roles in type I interferon response, such as *OAS2*, *OAS3*, *HERC5*, *IFIT2* and *IRF7*. Forty-five of 53 genes with higher expression in all tissues from FIP cats were identified as interferon stimulated genes (ISGs) in the Interferome v2.01 database (40) with particularly high expression of well-studied interferon response genes *ISG15*, *RSAD2*, *IFIT3* and *IFIT1* (Fig. 2b). The activation of type I interferon response in all tissues studied was confirmed by gene set enrichment analysis (GSEA) using Hallmark feline gene sets from MSigDB (41–43) (Fig. 2c). The other Hallmark feline gene set significantly enriched (FDR < 0.05) in all tissues from FIP cats was ‘Interferon gamma response’, which significantly overlaps with the ‘Interferon alpha response’ gene set. Several further Hallmark feline gene sets were significantly enriched in both liver and MLN, including ‘TNFα signaling via NFκB’, ‘inflammatory response’, ‘IL6 JAK STAT3 signaling’ and ‘Apoptosis’ (Fig. S2a). In addition, gene ontology (GO) enrichment analysis revealed higher expression of genes involved in B and T cell mediated immunity in multiple tissues from cats with FIP, with the strongest enrichment in MLN (Fig. 2d).

**Figure 2.**
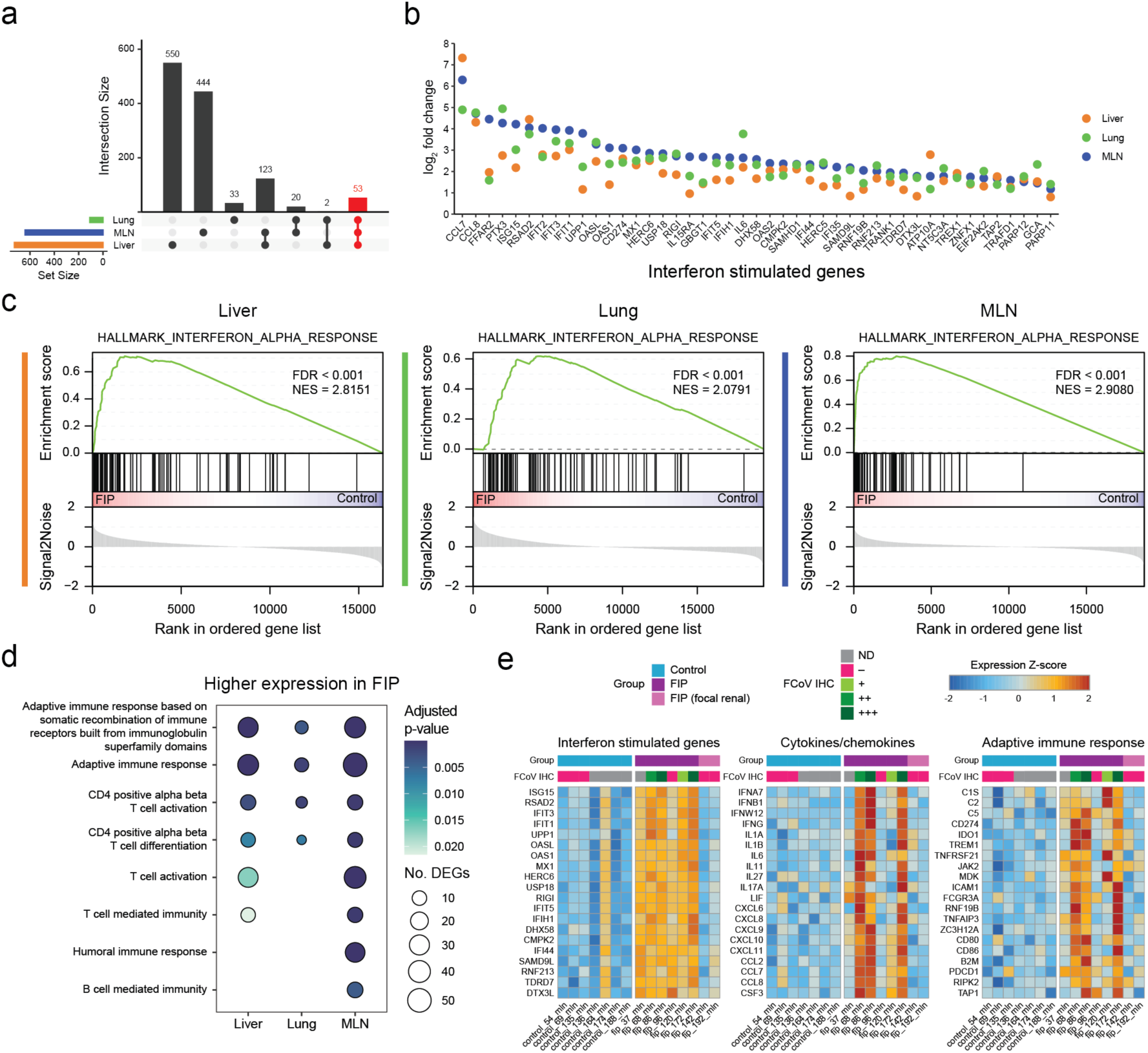
Identification of an immune response signature in multiple tissues from cats with FIP. (a) Upset plot showing overlap of differentially expressed genes with increased expression in FIP across all tissues. The set of genes with increased expression in FIP for all tissues is shown in red. (b) Plot of log2 fold changes for 45 interferon stimulated genes with higher expression in FIP in all tissues. For each gene, points represent the estimated fold change between FIP and control samples, with points coloured by tissue from which the sample was taken. (c) Gene set enrichment analysis (GSEA) plots for each tissue using the MSigDB Hallmark feline gene set ‘interferon alpha response’. FDR – false discovery rate, NES – normalised enrichment score. (d) Selected enriched gene ontology (GO) terms related to adaptive immune response for genes with higher expression in FIP in each tissue. Points are shown where each term was significantly (adjusted p-value < 0.05) enriched in each tissue, coloured by adjusted p-value, and the size of each point corresponds to the number of genes annotated with each term with significantly (adjusted p-value < 0.05) higher expression in FIP. (e) Heatmaps of scaled normalised expression for selected genes in all mesenteric lymph node samples, where rows are labelled with the gene for which expression is shown, and columns are labelled with the sample ID. Columns are annotated with sample group, and also the results of FCoV antigen detection by immunohistochemistry (IHC). ND – not determined.

Whilst DEGs involved in type I interferon response were consistently more highly expressed in the MLN of cats with FIP (but to a lesser degree in cats with focal renal FIP), DEGs encoding inflammatory cytokines/chemokines and proteins involved in the adaptive immune response were more heterogeneously expressed among cats with FIP (Fig. 2e, Fig S2b). These included *IL6* and *IL1B*, encoding proinflammatory cytokines previously found to be upregulated in cats with FIP (44), as well as *PDCD1* and *CD274*, encoding the immune checkpoint receptor PD-1 and its endogenous ligand PD-L1, respectively, which have previously been found to have higher expression in peripheral blood mononuclear cells (PBMCs) from cats with FIP compared to healthy controls (45). Inclusion of results of IHC for FCoV antigen suggested a correlation between virus abundance and inflammatory gene expression, indicating persistence of the virus despite pronounced immune activation (Fig. 2e).

### Assembly of full-length FCoV genomes from feline tissue RNA-seq samples

To further investigate the relationship between tissue FCoV abundance and gene expression, RNA-seq reads were re-mapped to a hybrid assembly comprising the feline reference genome (Felis_catus_9.0) and a previously published serotype I FCoV isolated from a cat with FIP (FCoV C1Je, DQ848678.1) (46). Reads originating from FCoV were detected at a medium-high level in at least one tissue from all cats with FIP, except for the two cats with renal FIP lesions, for which FCoV reads were not detected in the liver, lung or MLN (Fig. 3a). Notably, for one 16.2-year-old control cat (164), low abundance FCoV was detected in the liver, lung, and MLN, indicating systemic FCoV infection without development of FIP. Gene set scores were calculated for the DEGs encoding cytokines/chemokines and involved in adaptive immune response (from Fig. 2e), and linear regression analysis confirmed the correlation between viral load and expression of these gene sets across all tissues (Fig. 3b-c).

**Figure 3.**
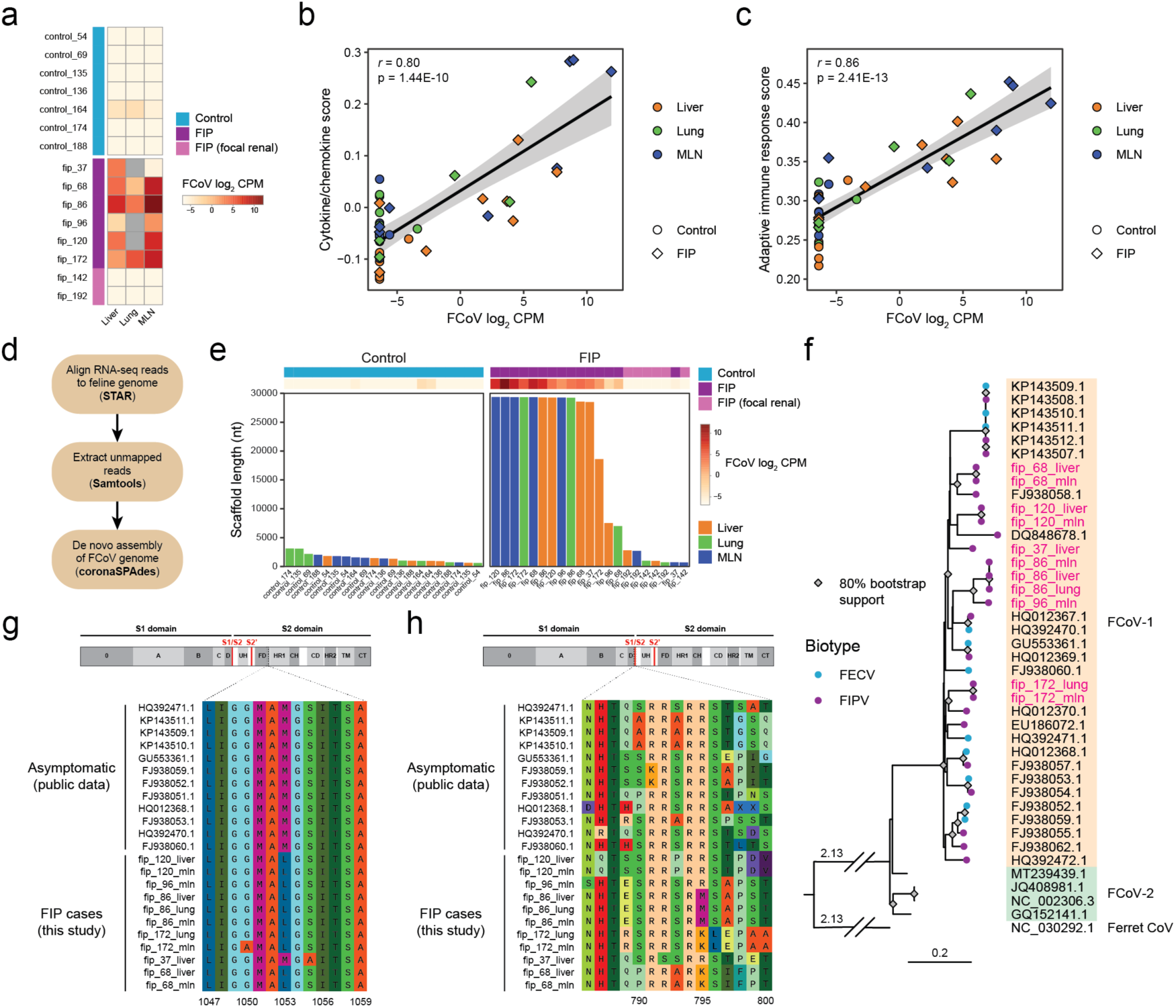
Assembly of full-length FCoV genomes from tissue RNA-seq of cats with FIP. (a) Heatmap showing normalised abundance (log2 counts per million mapped reads (CPM)) of RNA-seq reads mapped to the feline coronavirus (FCoV) genome (C1Je, accession DQ848678) in each sample. Grey cells indicate samples not available for inclusion in the study. (b) and (c) Correlation between FCoV normalised abundance and gene expression scores, calculated using normalised expression of genes shown in Fig. 2(e) encoding (b) cytokines and chemokines and (c) proteins involved in adaptive immune response. Each point represents an individual sample. The shape of each point corresponds to the sample group, and points are coloured by tissue from which the sample was taken. Regression lines are shown in black, with 95% confidence intervals shown in grey. Pearson’s r and corresponding p-values for each correlation are also shown. (d) Schematic overview of the analysis pipeline for assembly of feline coronavirus sequences from feline tissue RNA-seq reads. (e) Lengths in nucleotides (nt) of the longest scaffold assembled for each RNA-seq sample using coronaSPAdes. Bars are coloured by tissue of sample origin. Sample are labelled with FCoV normalised abundance, as shown in (a), and sample group. (f) Maximum likelihood tree based on whole genome alignment of FCoV sequences assembled in this study (sample names in magenta) and publicly available FCoV whole genomes (labelled with NCBI accession IDs). Branch tips are coloured by biotype, with published feline enteric coronavirus (FECV) sequences isolated from faeces of cats without clinical signs of FIP, and feline infectious peritonitis virus (FIPV) isolated from tissues or effusions of cats with FIP. Clades are coloured and labelled with the corresponding FCoV serotype: serotype 1 (orange) or serotype 2 (green). Nodes with >80% bootstrap support are indicated with grey diamonds. A ferret coronavirus genome (accession NC_030292.1) was used to root the tree. Branch lengths are proportional to the number of nucleotide substitutions per site. (g) and (h) Multiple sequence alignments of predicted Spike protein sequences from public FECV genomes and the genomes assembled from cats with FIP as part of this study, showing mutations in (g) the S1/S2 cleavage site and (h) the ‘M1058L’ substitution in the putative fusion domain used clinically for diagnosis of FIP. For each alignment, a schematic representation of the FCoV Spike protein domain structure (from PDB 6JX7) is shown: UH – upper helix, FD – fusion domain, HR1 – heptad repeat 1, CH – central helix, CD – connector domain, HR2 – heptad repeat 2, TM – transmembrane domain, CT – cytoplasmic tail. Functionally important cleavage sites (S1/S2 and S2’) are shown in red.

Recovery of FCoV genomic sequence from feline tissue RNA-seq samples was attempted by metagenomic assembly of the unmapped reads following alignment to the feline reference genome (Fig. 3d). For 11/25 samples from FIP cats, the largest assembled scaffold was approximately the full length of the FCoV genome (∼29 kb), and shorter scaffolds containing FCoV sequence were assembled in a further 3 FIP samples (Fig. 3e). In all remaining samples from FIP and control cats, scaffolds <5 kb were subjected to a BLAST search and found to contain sequences of feline rather than viral origin. Phylogenetic analysis revealed that all assembled FCoV sequences clustered with public FCoV-1 sequences, with public FCoV-2 genomes forming a distinct clade. Additionally, in agreement with previous observations (47), there was no clear separation of FCoV genomes identified as FIPV or FECV, providing further support for a lack of clearly discriminating genetic signal between the two biotypes (Fig. 3f).

As expected, previously identified virus mutations associated with systemic spread were detected in the FCoV assemblies generated in this study, such as the M1058L and S1060A mutations in the S gene (36) (Fig. 3h). Diverse mutations in the putative S1/S2 cleavage site, which have also been associated with FIP (37), were detected in the RNA-seq FCoV genome assemblies (Fig. 3g). Several of the assembled FCoV genomes also contained truncations of ORF3c, which has been only partially associated with FIP, as some FIPV isolates retain full-length ORF3c, as was the case in this study (Fig. S3a). Other notable features of the assembled FCoV genomes included large deletions in the N-terminal domain 0 of the predicted Spike protein for two assemblies, both from liver samples, for FIP cats 37 and 68 (Fig. S3b). In the case of cat 68, the FCoV genome assembled for the MLN sample contained a full-length S gene, indicating that this mutation had been acquired within the host (Fig. S3c).

These results demonstrate the feasibility of recovering full-length FCoV genomes from RNA-seq of tissues from cats with FIP. The FCoVs detected in this study were all FCoV-1 and their genomes contained features previously associated with FIP or systemic spread of the virus.

### Analysis of FCoV intrahost single nucleotide variants (iSNVs)

Although feline tissue RNA-seq reads that were not mapped to the feline genome aligned to publicly available FCoV genomes, coverage was relatively poor, particularly for the S gene region encoding the N-terminal domain of the spike protein (Fig. S4a). S gene pairwise nucleotide identities between FCoV assemblies from different cats were in some cases as low as ∼82%, with significantly higher (>98%) identity between assemblies from different tissues from the same cat (Fig. S4b). To overcome this, for each individual cat the RNA-seq reads that did not map to the feline genome from different tissues were aligned to the FCoV genome assembled from MLN RNA-seq reads for the same cat. The MLN assemblies were chosen as reference sequences to enable within-host analysis of FCoV genetic variation because the MLN is the presumed first site of infection after FCoV spreads from the gut (48), and also because, for most of the FIP cats with detectable FCoV in the liver, lung or MLN, the mean depth of RNA-seq reads mapped to the respective FCoV assembly was highest for the MLN sample (Fig. S4c). For further analysis of iSNVs, samples from FIP cats 37, 142, and 192 were excluded as full-length FCoV genome assemblies were not obtained using RNA-seq reads from the MLN of these cats.

After filtering, a total of 1,344 SNVs were identified for all remaining samples with respect to the MLN FCoV assembly from the same cat, and the number of variants detected in each sample was dependent on the mean depth of reads mapped to the respective MLN assembly (Fig. 4a). The full list of quality-filtered variants detected in all tissues is available in Supplementary Table S6. Downsampling the reads from the four samples with the highest mean depth (MLN from FIP cats 120, 172, 68 and 86) to fixed values revealed that detection of variants was stable, relative to calls from the maximum number of available reads, at and above 500x (Fig. S4d). None of the available tissue samples for FIP cat 96 exceeded this threshold, and these samples were therefore excluded from further analysis. In the remaining cats, annotation of detected variants revealed non-synonymous mutations in all of the annotated ORFs in the assembled FCoV genomes, with a particularly large number of mutations in samples from FIP cat 172 (Fig. 4b). Inspection of the alternate allele frequency (AAF) distribution for variants detected in the MLN sample from cat 172 revealed a high frequency of variants with an AAF of ∼0.20 (Fig. S4e). This was not seen for variants detected in the MLN of cats 120, 68, and 86. One possible explanation is that this represents co-infection with at least two strains of FCoV in cat 172, with the major and minor strains present in the MLN at a ratio of ∼4:1.

**Figure 4.**
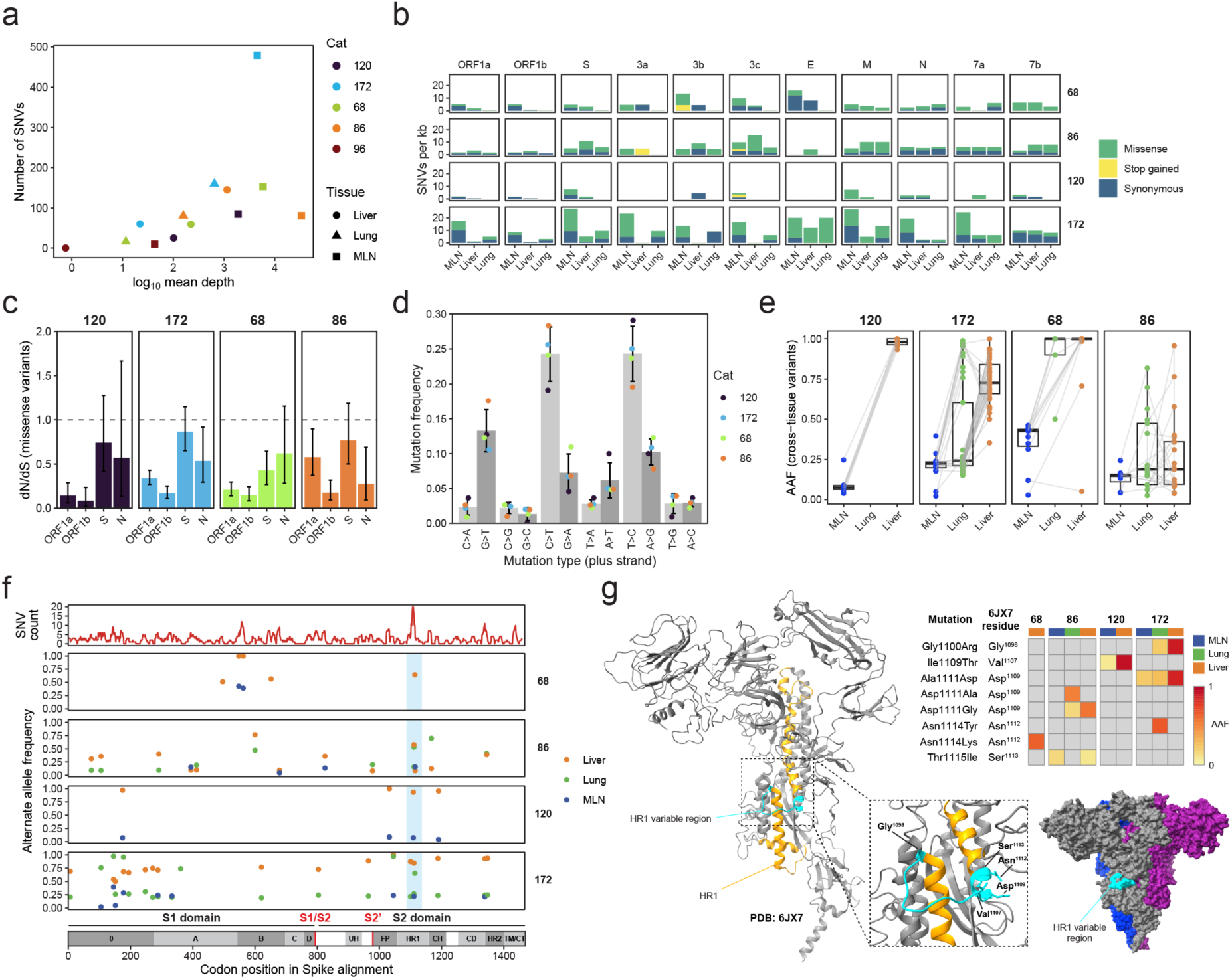
Analysis of FCoV intrahost genetic variation. (a) Plot of log_10_ mean depth of RNA-seq reads from feline tissues aligned to the FCoV assembly from the respective cat’s MLN sample versus the number of single nucleotide variants (SNVs) called with LoFreq. Samples from FIP cats 37, 142 and 192 were not included due to the lack of a full-length FCoV genome assembly from the MLN of these cats. Each point represents an individual sample. The shape of each point corresponds to the tissue from which the reads were obtained, and points are coloured by FIP cat from which the sample was obtained. (b) Counts of FCoV SNVs of different types, normalised to SNVs per kilobase (kb), called for each tissue sampled from the FIP cats with at least one tissue with > 500x mean depth of aligned reads (rows), and further divided by FCoV ORF (columns). (c) Maximum likelihood estimates of dN/dS for non-synonymous substitutions in genes reaching significance (Benjamini-Hochberg adjusted p-value < 0.1) in at least one cat. The dashed horizontal line at dN/dS = 1 marks the value expected under neutral selection. Identical non-synonymous variants detected in multiple tissues from the same cat were deduplicated to avoid biasing the dN/dS estimates. Error bars show the 95% confidence intervals of maximum likelihood estimates, which are represented by the height of the bars. (d) Mutation spectrum of all SNVs detected across all tissue samples from cats with > 500x mean depth in at least one sample. Points represent the frequency of each mutation in all tissue sample from each cat. Bar heights represent the mean frequency for each mutation across all cats, and error bars represent the standard deviation. Mutations are grouped in pairs corresponding to equivalent mutations occurring on either the plus (light grey) or minus (dark grey) strand FCoV RNA. (e) Alternate allele frequencies (AAF) of SNVs detected in multiple tissues from the same cat. Each point represents an individual SNV, and lines connect identical SNVs (by position and alternate allele) detected in different tissues from the same cat. Upper and lower box edges represent the third and first quartiles, respectively, with the centre line indicating the median value. Whiskers extend to the maximum (upper) or minimum (lower) value that is no more than 1.5 x the interquartile range from the respective box edge. (f) Focused view of non-synonymous SNVs detected in multiple tissues, or at > 0.5 AAF in the liver or lung, in the S gene. The top track (red line) shows a sliding window analysis (window size = 10 residues) of the density of all SNVs detected in all samples, with adjusted codon positions from a multiple sequence alignment of predicted spike protein sequences from the MLN FCoV assemblies of the cats shown. Each point represents an individual SNV, and points are coloured by tissue in which the SNV was detected. A variable region common to all cats, with high SNV density, is highlighted in light blue. The bottom track shows the domain structure of FCoV spike protein, with residue numbers derived from a previously published cryo-EM structure (PDB: 6JX7) and adjusted based on alignment with the predicted spike protein sequences generated in this study. UH – upper helix, FP – fusion peptide, HR1 – heptad repeat 1, CH – central helix, CD – connector domain, HR2 – heptad repeat 2, TM/CT – transmembrane/cytoplasmic tail. (g) Left - structural model of the FCoV spike monomer (PDB: 6JX7) with the HR1 domain coloured orange, and the variable region identified in panel (f) coloured cyan. The box contains a zoomed-in view of the variable region, with the residues corresponding to mutations in the FIP cats from this study labelled. Bottom right – surface visualisation of the spike trimer (PDB: 6JX7) with different colours for each monomer, and the variable region in HR1 coloured cyan on the protein surface. Top right – heatmap showing the AAFs of mutations detected in each tissue from each FIP cat, along with the corresponding residue in the cryo-EM structure.

When comparing the relative frequency of synonymous and non-synonymous mutations, at the level of the whole FCoV genome, negative selection was detected for non-synonymous mutations in all cats analysed, with strong purifying selection detected for nonsense variants (Fig. S4f). Gene-level estimates of selection pressure were restricted by the limited number of variants detected in shorter ORFs in each cat, however significant purifying selection was detected in at least one cat for the ORF1a, ORF1b, S and N genes (Fig. 4c). In contrast, dN/dS estimates > 1, indicating positive selection, were obtained in multiple cats for M and ORF7b, however the limited number of variants detected in these genes resulted in wide confidence intervals, and dN/dS > 1 for these genes was not obtained in all FIP cats included in the analysis (Fig. S4g). Analysis of the specific nucleotide substitutions detected in FCoV from each cat revealed a consistent signature of over-represented mutations, with strong imbalance between equivalent mutations on the plus and minus strands of FCoV RNA (Fig. 4d). Multiple host factors are known to induce mutations in viral RNA, including the protein encoded by the interferon stimulated gene *ADAR* and APOBEC family proteins. Here, *ADAR* had significantly (adjusted p-value < 0.05) higher expression in the liver and MLN of FIP cats compared to control cats (Fig. S4h), however *APOBEC1* expression was not detected in any tissue studied and other annotated feline APOBEC family genes were not differentially expressed between FIP and control cats. The trinucleotide sequence context of C>T mutations showed a preference for substitutions at sites flanked by A or T nucleotides, indicative of APOBEC-mediated editing, but there was no clear preference in the sequence context of T>C mutations (Fig. S4i).

Of the 1,344 SNVs detected across all samples, 182 (13.54%) were shared SNVs detected in multiple tissues from the same cat, representing 82 distinct sites. Comparison of shared SNV frequencies among tissues indicated that, while most SNVs had increased AAFs in the liver or the lung compared to the MLN, some decreased in frequency. There was no consistent pattern in AAF changes between tissues (Fig. 4e). In the case of FIP cat 172, which had already been identified as a possible case of co-infection with a mixture of FCoV strains, the MLN minor variants at an AAF of ∼0.2 were split, with some remaining at a similar AAF, and others approaching fixation of the minor allele. This could be explained by the presence of more than one minor strain in the original MLN sample, suggesting even more complexity in the FCoV genetic factors influencing disease. Focused analysis of non-synonymous SNVs in the S gene revealed largely distinct patterns of mutation among cats. However, a ∼45 nt site within the region encoding the heptad repeat 1 (HR1) domain of the spike protein contained at least one cross-tissue non-synonymous SNV, or a non-synonymous SNV with AAF > 0.5 in either the liver or lung, in all four cats analysed (Fig. 4f). Alignment of these mutations with a previously published cryo-EM structure of FIPV spike protein (PDB: 6JX7) predicts that they alter the sequence of an exposed loop between alpha helices of the HR1 domain (Fig. 4g). As well as the S gene, non-synonymous mutations across multiple tissues from all four cats were concentrated in a specific region of the M gene, likely encoding a domain expressed on the surface of the virion, and hence possibly susceptible to host immune recognition (Fig. S4j).

### The T cell receptor (TCR) repertoire in FIP

The TCR repertoire shapes the adaptive immune response, and we hypothesised that differences would exist in the immune repertoire between FIP and control cats, potentially providing insight into the immune dysregulation and hyperglobulinaemia in FIP. Due to the presence of reads mapping to regions encoding TCRs in the RNA-seq analysis, initially, TCR α and β chain clonotypes were assembled from RNA-seq reads from all samples. As expected, due to the higher expression of immune response genes and predicted immune activity of the tissue, the number of assembled clonotypes was highest in MLN samples (Fig. S5a-b).

Preliminary analysis identified sharing of clonotypes among FIP samples (Fig. S5c-d). In order to better characterise the TCR repertoire in FIP, targeted TCR sequencing was performed on a larger cohort of feline MLN samples in the same archive, from cats with and without FIP, including the MLN samples subjected to RNA-seq. Targeted TCR sequencing libraries were filtered for low read counts, resulting in repertoires for analysis from 19 FIP cats and 22 control cats. Targeted sequencing identified a greater number of TCR clonotypes than RNA-seq in all samples analysed with both methods (Fig. S5e), and for TCRα clonotypes detected by both RNA-seq and targeted TCR sequencing the proportion of reads assigned to top clonotypes across samples was highly correlated (Fig. S5f). When grouped by presence/absence of FCoV antigen as determined by IHC, there was a significant decrease in repertoire diversity (measured using the Chao1 index for species richness) in FIP FCoV+ samples compared to both FIP FCoV- and control samples for both the α and β chain (Fig. 5a). Unevenness in the distribution of sequencing reads across clonotypes (measured by Gini coefficient) was significantly increased in FIP FCoV+ samples compared to both FIP FCoV- and control samples for both chains (Fig. 5b). This implies relatively increased clonal expansion, and therefore T cell activation, in the FIP MLN samples positive for FCoV. This phenomenon was also apparent in a sample-level analysis of clonotype abundance as a proportion of repertoire space, with a greater proportion of the TCR α and β repertoires in FIP FCoV+ samples occupied by clonotypes with high counts compared to FIP FCoV- or control samples (Fig. 5c).

**Figure 5.**
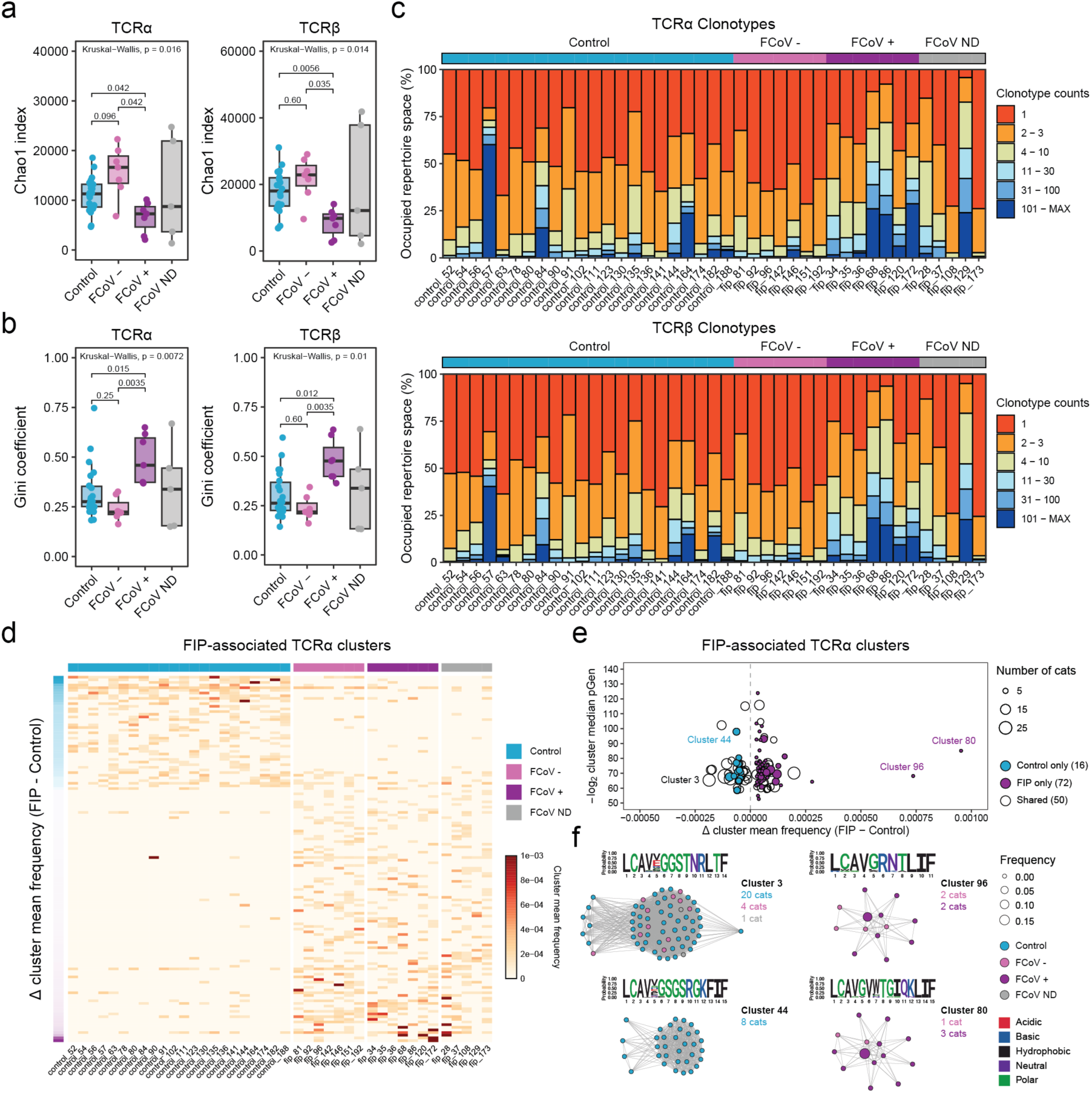
Identification of FIP-associated T cell receptor (TCR) clonotypes. (a) and (b) Boxplots showing (a) Chao1 index and (b) Gini coefficient values for TCRα and TCRβ repertoires assembled from targeted TCR sequencing of mesenteric lymph node samples from cats with and without FIP. For estimation of each diversity index, repertoires were downsampled to the number of reads in the repertoire with the lowest sequencing depth. Each point represents an individual sample, and points are coloured by sample group. FIP samples are divided into groups according to positivity for FCoV antigen by immunohistochemistry (IHC). FCoV ND – not determined. Upper and lower box edges represent the third and first quartiles, respectively, with the centre line indicating the median value. Whiskers extend to the maximum (upper) or minimum (lower) value that is no more than 1.5 x the interquartile range from the respective box edge. Groups were compared using the Kruskal-Wallis test, followed by pairwise Wilcoxon tests with correction for multiple testing using the Holm method. (c) Proportion of each individual TCRα (top) and TCRβ (bottom) repertoire occupied by clonotypes with specified counts. Samples were grouped by sample status, with FIP samples grouped by FCoV positivity by IHC. (d) Heatmap showing, in each sample (columns), the mean frequency of clonotypes contained in FIP-associated TCRα clusters (rows). Clusters are ordered by the difference in cluster mean frequency between FIP and Control samples. Samples are grouped by sample status, with FIP samples grouped by FCoV positivity by IHC. (e) Plot showing the relationship between difference in cluster mean frequency between FIP and Control samples and the median cluster generation probability (pGen). Each point represents an individual TCRα cluster, and the size of each point is proportional to the number of cats with clonotypes included in the cluster. Filled points are exclusive to FIP (purple) or Control (blue) cats, and unfilled points represent clusters comprised of clonotypes from both FIP and Control cats. (f) Selected FIP-associated TCRα clusters common to a large number of cats (Cluster 3), with markedly low pGen (Cluster 44) or with extreme differences in cluster mean frequency between FIP and Control (Clusters 96 and 80). Nodes represent individual TCRα clonotypes, and edges connect clonotypes where the CDR3 sequence differs by a single amino acid. Nodes are coloured by sample group of the cat in which the clonotype was sequenced, and node size is proportional to the frequency of the clonotype in its original repertoire. Sequence logos represent the CDR3 sequences of all TCRα clonotypes in each cluster, with amino acids coloured by side chain chemistry. For each cluster shown, the number of cats from which constituent clonotypes were assembled is shown, coloured by sample group.

To identify TCR clonotypes associated with FIP, network analysis was performed using all available clonotype data to identify clusters with a maximum Hamming distance of 1. Previous work has shown that, in humans, such clonotypes are likely to bind identical or highly similar peptide-MHC complexes (49,50). CDR3 sequences significantly associated with either FIP or control groups were identified (p < 0.05, Fisher’s exact test) and TCR clonotypes with highly similar CDR3 sequences across all samples were used to build separate networks for the α (Fig. S5g) and β (Fig. S5h) chains. One hundred and thirty-eight TCRα clusters with FIP-associated CDR3 sequences were identified, with either higher or lower mean cluster frequency in FIP samples compared to controls (Fig. 5d). A smaller number of TCRβ clonotypes were found to be associated with FIP, with limited sharing of clonotypes among either FIP or control samples (Fig. S5i). Generation probabilities (pGen), which reflect the likelihood of a CDR3 sequence being generated in any individual through V(D)J recombination, were calculated for the CDR3 sequences of FIP-associated clonotypes in each cluster, revealing that the majority of FIP-specific and control-specific clusters had a relatively high median pGen (Fig. 5e, Fig. S5j). A small number of clusters, however, had relatively low median pGen, indicating that differences in prevalence and frequency of these clusters between FIP and control samples are more likely to be driven by selection following antigenic stimulation of T cells. Clusters with higher mean frequency in FIP samples were generally marked by very high frequency in a single sample, with limited sharing of similar clonotypes among samples. Clusters with higher mean frequency in control samples, however, were more consistently represented across the majority of control samples and were largely absent from FIP samples (Fig. 5f). These results suggest that, in cats with FIP, the aberrant T cell response is directed by multiple diverse epitopes, and that the majority of the associated clonotypes result in an ineffective immune response to FCoV, given the persistence of virus in these samples. Importantly, the presence of clonotypes highly represented among control samples and virtually absent from FIP samples also suggests the possibility of protective TCR clonotypes.

### The B-cell receptor (BCR) repertoire in FIP

Exploration of the BCR repertoire in FIP is particularly valuable given the hyperglobulinaemia associated with the clinical presentation, however the *Felis catus* immunoglobulin heavy chain (IGH) locus remains uncharacterised in the IMGT database (51), limiting the feasibility of feline BCR repertoire analysis. To address this, a combination of *de novo* VDJ gene prediction and manual annotation was used to map the feline IGH locus and construct a reference sequence library for BCR heavy chain sequence analysis (Fig. S6a) (see Methods). Using this library, IGH clonotypes were assembled from all RNA-seq samples, and analysis of clonotype abundances revealed greater expansion of clonotypes in FIP samples compared to controls, particularly in the MLN (Fig. 6a). In FIP samples, clonal expansion (measured by Gini coefficient) was significantly correlated with FCoV normalised abundance in MLN samples, but not in liver or lung (Fig. 6b), likely reflecting the absence of germinal centres found in secondary lymphoid tissues.

**Figure 6.**
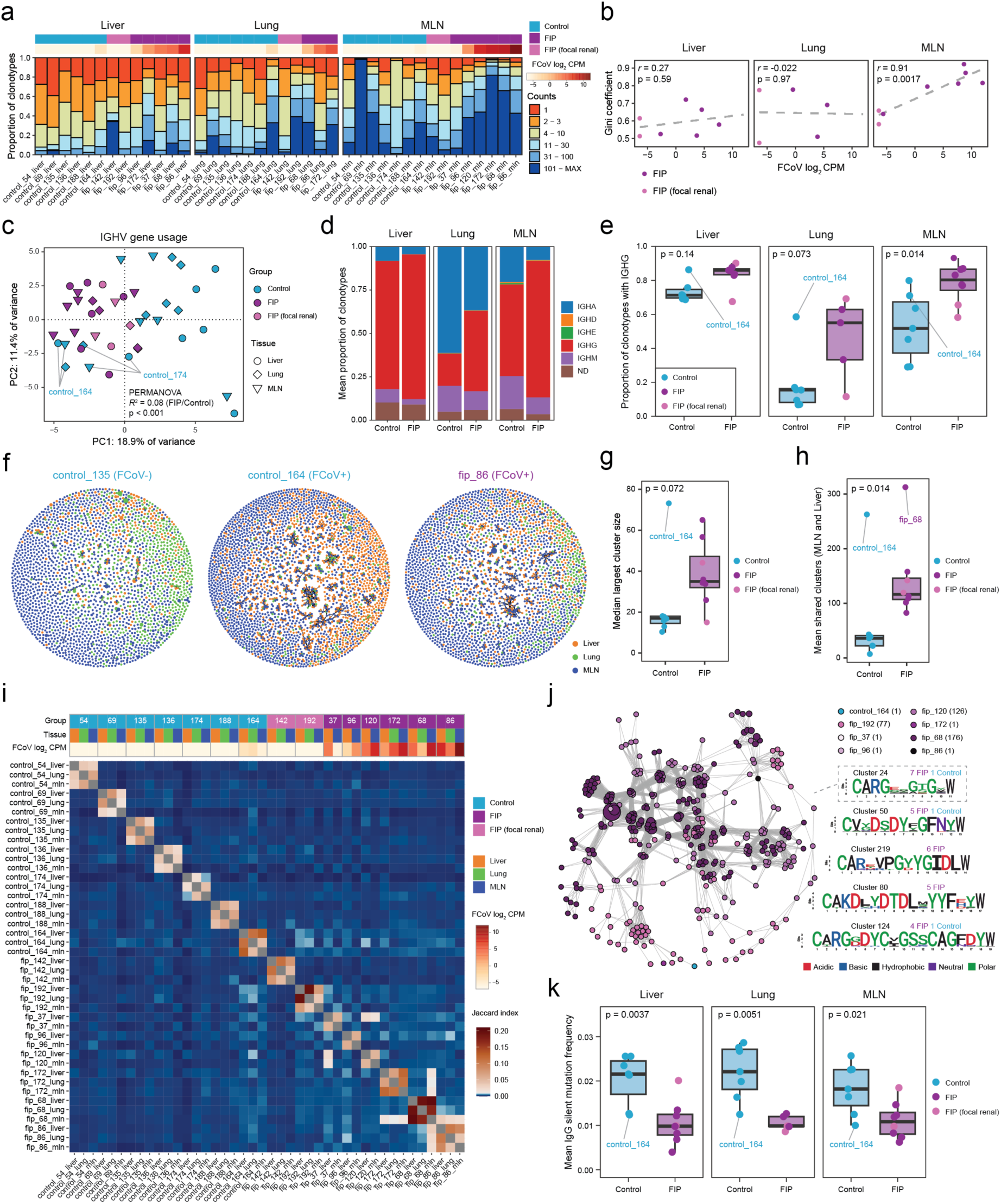
Changes in the immunoglobulin heavy chain repertoire associated with FIP. (a) Proportion of immunoglobulin heavy chain (IGH) clonotypes in each RNA-seq sample with specified counts. Samples are annotated with sample group and FCoV normalised abundance. (b) Plots showing relationship between FCoV normalised abundance and Gini coefficient for IGH clonotypes in each tissue from cats with FIP. Each point represents an individual sample, and points are coloured by sample group. Dashed grey lines indicate results of linear regression, and Pearson’s r and associated p-value is shown for each correlation. (c) Results of principal component analysis (PCA) based on IGHV gene usage in IGH repertoires from all RNA-seq samples. Each point represents an individual sample, with point shape corresponding to tissue, and colour corresponding to sample group. Results of PERMANOVA test for difference in IGHV usage between FIP and control are shown, where cat age was included as a covariate. (d) Mean proportion of clonotypes with the indicated constant region in FIP and Control samples for each tissue. ND – not determined. (e) Boxplots showing the proportion of clonotypes in FIP and Control samples, in each tissue, with the constant region identified as IGHG. Each point represents an individual sample, and points are coloured by sample group. FIP and Control samples were compared using the Wilcoxon rank sum test. (f) Representative IGH repertoire networks constructed using clonotypes from all tissues for selected cats. Nodes represent clonotypes with unique CDR3 nucleotide sequences, and edges connect sequences that differ by ≤ 1 nucleotide. Nodes are coloured by tissue from which the clonotype was assembled, and node size is proportional to the frequency of the clonotype in the original repertoire. (g) and (h) Boxplots showing the (g) median largest cluster size and (h) mean number of clusters shared by MLN and liver samples for subsampled repertoire networks from each cat. Each point represents an individual cat, and points are coloured by sample group. (i) Matrix of overlap values (Jaccard index) between RNA-seq samples for clusters in a global network comprising all IGH clonotypes from all samples. Clonotypes were clustered by CDR3 sequence, with nodes representing unique CDR3 sequences, and edges connecting sequences that differ by ≤ 1 amino acid. Samples are annotated with cat ID, sample group, tissue and FCoV normalised abundance. (j) Network representation of cluster 24 from the global IGH repertoire network analysis, comprised of clonotypes from 7/8 FIP cats and a single clonotype from 1 Control cat (control_164). Nodes represent individual clonotypes, and edges connect clonotypes with CDR3 sequences that differ by ≤ 1 amino acid. Nodes are coloured by cat from which the clonotype was assembled, and node size is proportional to the frequency of the clonotype in its original repertoire. For each cat, the number of nodes contained in cluster 24 is shown in brackets. Also shown are amino acid sequence logos for the CDR3 of the top 5 clusters by number of FIP cats included, with cluster 24 enclosed in a dashed grey box. For each sequence logo, the number of FIP and Control cats included in the cluster is shown, and amino acids are coloured by side chain chemistry. (k) Boxplots showing the mean silent (synonymous) mutation frequency in full-length IgG clonotypes assembled for FIP and Control RNA-seq samples from each tissue. Each point represents an individual sample, and points are coloured by sample group. FIP and Control samples were compared using the Wilcoxon rank sum test. For all boxplots, upper and lower box edges represent the third and first quartiles, respectively, with the centre line indicating the median value. Whiskers extend to the maximum (upper) or minimum (lower) value that is no more than 1.5 x the interquartile range from the respective box edge.

Although limited differences in specific IGHV gene usage were detected between FIP and control samples, with only minor concordance among tissues (Fig. S6b), principal component analysis of IGHV gene usage in all tissue samples revealed a significant difference between FIP and control samples (p < 0.001, PERMANOVA) (Fig. 6c). Tissues from 2 control cats appeared more similar to FIP samples than other control samples: control_164, which was found to be FCoV-positive by RNA-seq with a histological diagnosis of metastatic haemangiosarcoma, and control_174, an 8-month-old cat with a diagnosis of portovascular shunt and hepatic encephalopathy but no evidence of FIP or systemic FCoV infection. Analysis of IGH constant gene usage revealed differences in isotype frequency among tissues, and a higher mean proportion of clonotypes using IGHG was detected in all 3 tissues from FIP cats compared to controls (Fig. 6d). The difference in IGHG usage was significant only in the MLN, due to a marked increase in IGHG usage in the liver and lung samples from control_164 (Fig. 6e). These differences were consistent with FCoV-specific shaping of the IGH repertoire, and with class switch recombination driving increased IgG production in FIP cats.

To assess IGH repertoire diversity across tissues, individual repertoire networks were constructed for each cat using all clonotypes from all tissues, and then repeatedly subsampled to produce representative visualisations and network statistics (see Methods). Representative networks are shown for selected samples in Fig. 6f, and for all samples in Fig. S6c. The repertoire for control_135 was largely comprised of individual, disconnected nodes (clonotypes), whereas networks for control_164 (with systemic FCoV, but without FIP) and fip_86 revealed the presence of large clusters of connected nodes (CDR3 nucleotide Hamming distance ≤ 1). The latter represents expansion and diversification of IGH sequences, likely through somatic hypermutation (SHM) following antigen-specific B-cell activation. The median largest cluster size for subsampled repertoire networks was higher in FIP samples compared to controls, with the exception of control_164, which had the highest value of all samples (Fig. 6g). The mean number of clusters containing clonotypes found in both the liver and MLN was compared between FIP and control samples (lung was excluded due to some FIP cats without lung samples included in the study) and was significantly higher for FIP samples compared to controls (Fig. 6h), again with the exception of control_164, indicating a greater number of circulating clonal lineages of B cells specific for similar antigens in FIP cats.

To identify convergent clonotypes between cats, a global network was constructed using all clonotypes from all samples (CDR3 amino acid Hamming distance ≤ 1). Membership of the resulting clusters was compared across all samples, revealing strong overlap between different tissues of the same cat, but weak overlap among the same tissues from different cats (Fig. 6i). The degree of within-cat overlap was higher for FIP samples than for controls (Fig. S6d). Between-cat overlap was limited in general, but was higher in FIP samples, indicating the presence of rare convergent B-cell responses in cats with FIP. The global network cluster containing clonotypes from the largest number of FIP cats included 7/8 FIP cats (excluding only fip_142, a case with exclusively renal FIP lesions) (Fig. 6j). The top 5 clusters, by number of FIP cats represented, had diverse CDR3 sequences, with variable lengths and amino acid composition, reflecting the polyclonal BCR response against, presumably, diverse epitopes of FCoV in FIP.

Quantification of the frequency of somatic hypermutation in assembled full-length BCRs revealed a lower frequency of silent mutations (frequency of which is not influenced by selection based on affinity for specific antigens but may influence transcription and translation efficiency) in the variable region of IgG clonotypes in all tissues from cats with FIP (Fig. 6k). This was also seen for other isotypes, with a higher density of low or zero mutation variable region sequences in FIP samples compared to controls (Fig. S6e). Analysis of selection scores, however, revealed strong positive selection in complementarity determining regions (CDRs) of IGH clonotypes from FIP cats, with neutral selection scores for control cats (Fig. S6f). In framework regions (FRs), there was strong negative selection for both FIP and control samples. This could be explained by recent expansion of nascent clonotypes relatively unaltered by SHM in FIP samples, compared to longer-lived clones that have diverged from germline sequences in control samples. Given the significant difference in the ages of FIP and control cats, an age effect could not be ruled out, however within each group (FIP and control) there was no significant correlation between age and mean mutation frequency (Fig. S6g).

Taken together, these results demonstrate that the IGH repertoire of cats is substantially altered across multiple tissues by the presence of FCoV, and that convergent B-cell responses, although rare, do occur in FIP.

## Discussion

This is the first study to show that expression of proinflammatory cytokines and adaptive immune response genes are highly correlated with FCoV abundance across tissues in cats with FIP. The persistence of viral RNA despite immune activation and proinflammatory signaling provides clear evidence for the failure of the feline immune system to restrict viral replication in many cats with FIP. It also explains why direct-acting anti-viral therapy (i.e. inhibiting viral replication) is necessary in the successful treatment of FIP (3–5). This is in contrast to the use of antivirals in severe COVID-19 cases in humans, where more modest impacts on disease progression are seen (52).

Type I interferon signaling is activated in a wide range of viral infections, inducing expression of numerous ISGs with antiviral and immunomodulatory functions. In other coronavirus infections, such as SARS-CoV and MERS-CoV in humans, dysregulated interferon signaling coupled with robust virus replication results in hyperinflammation and more severe disease (53). It is therefore not surprising that inborn errors in interferon signaling, including polymorphisms in ISGs and interferon autoantibodies, have been associated with severe COVID-19 in humans (54). Furthermore, SARS-CoV-2 proteins have been shown to antagonise ISG expression (55,56). In the present study, activation of type I interferon signaling was detected in the liver, lung and MLN from FIP cats, as well as an increase in expression of pro-inflammatory cytokines and genes involved in the adaptive immune response in samples with high abundance of FCoV RNA.

Some cytokines were significantly more highly expressed in FIP cats across all 3 tissues, although their expression was heterogeneous amongst samples and correlated with FCoV abundance. This included *IL6*, previously associated with FIP and also linked to disease severity in COVID-19 (44,57,58). In contrast, the MLN appeared to be the main tissue focus of increased expression of other proinflammatory cytokines (e.g. *IL1B* and *CXCL8*). Current understanding of the T cell response to FCoV is limited, although apoptotic T cell depletion is common in cats with FIP (59,60), and production of IFN-γ primarily by activated T cells is reportedly correlated with FIP progression (61). In this study, higher expression of *IFNG* in the MLN of FIP cats compared to controls was significant (adjusted p-value < 0.05), and nominally significant in the liver and lung (raw p-value < 0.05). This is consistent with previous studies that detected high expression of IFN-γ in lesions and effusions from FIP cats (48,62), and suggests a complex role for IFN-γ in the response to FCoV, as high expression in PBMCs upon exposure to FCoV has previously been associated with protection from FIP, likely due to early and effective induction of systemic cell-mediated immunity (61,63).

Of further note, and potential future therapeutic significance, higher expression of *CD274* encoding PD-L1, a ligand for the immune checkpoint receptor PD-1 (*PDCD1*), was also detected in all 3 tissues from cats with FIP. Coronaviruses, such as SARS-CoV-2, can stimulate increased expression of PD-L1 and thereby promote T cell ‘exhaustion’ (64,65). As a potent inhibitor of antigen-specific T cell responses, blocking the interaction of PD-1 and its ligands (including PD-L1) has been explored as a potential therapy in human infectious diseases after showing promise in cancer immunotherapy (66). An anti-feline PD-1 antibody has recently been developed and can block the interaction of PD-1 and PD-L1 *in vitro* (67). Similar to several inflammatory markers, *CD274* expression was significantly correlated with FCoV abundance (all tissue samples, r = 0.68, p = 8.66 x 10^-8^), driven mainly by expression in MLN and liver. This complements the findings of a previous study where *PDCD1* and *CD274* were more highly expressed in PBMCs from FIP cats compared to healthy controls (45).

Most FCoV infections only affect the gut and resolve naturally, although some cats continue to shed virus intermittently (68). FIP was historically presumed to be an inevitable consequence of systemic FCoV infection. However, reported cases of FCoV infecting tissues other than the gut in cats without clinical FIP indicate that systemic infection alone is not sufficient to cause FIP (69). This is illustrated in the present study by the detection of FCoV reads in all 3 tissues of one 16.2-year-old control cat (control 164) with similar levels of ISG expression to FIP cats. Importantly this cat, diagnosed with metastatic haemangiosarcoma, did not demonstrate a tissue signature of proinflammatory and adaptive immune response gene activation.

The present study has also provided an opportunity to examine mutations in FCoV sequences found outside the feline gut. While FCoV mutations within the host are still considered likely to play a role in altering the virus’ capacity to infect monocytes/macrophages, these mutations alone are not enough to induce FIP (69). As expected, previously described mutations were detected in the FCoV genomes assembled from FIP cats in this study, supporting their association with systemic spread beyond the gastrointestinal tract, however there was no single mutation common to all isolates. There was evidence of within-host FCoV mutation, as in FIP cat 68 ORF3c was truncated in the liver but intact in the MLN FCoV assembly. Similarly, in FIP cat 86, with intact ORF3c present in all FCoV assemblies, a frameshift deletion was detected (AAF = 0.23) in the liver sample, revealing a minor population of FCoV with truncated ORF3c. Large in-frame deletions in domain 0 of Spike are a characteristic feature of the novel, highly pathogenic recombinant FCoV-23 causing a recent FIP outbreak in Cyprus (28) and have been shown to increase FCoV-23 fusogenicity *in vitro* (70), but are found only rarely in FCoV-1 isolates (71). In the present study, there was evidence of within-host acquisition of such a mutation between the MLN and liver of FIP cat 68, but the significance of these rare deletions in FCoV-1 remains unknown, particularly as the host cell entry receptor for FCoV-1 has yet to be identified but is distinct from that of FCoV-23 and other FCoV-2 strains (70,72).

This multi-tissue study also enabled genome-wide evaluation of FCoV iSNVs, which are an important source of emergence of potentially harmful variants. The mutational signature of FCoV is similar to that of SARS-CoV, SARS-CoV-2 and MERS-CoV in humans, suggesting common influences on coronavirus evolution between species (73). These include the double-stranded RNA-specific deaminase ADAR, which was more highly expressed in tissues from FIP cats compared to controls, and accumulated mutations in the minus strand of the transitory duplex formed during FCoV genome replication were likely detected as T>C mutations in the more abundant plus strand. Detection of a high frequency of C>T FCoV mutations implies a role for APOBEC-mediated RNA editing of the FCoV genome, with APOBEC1 being the best-characterised RNA-editing enzyme in this family. Plus strand RNA, including FCoV genomic RNA and subgenomic RNA (sgRNA), is expected to be more abundant at any one time, allowing APOBEC-mediated C>T mutations to accumulate. Importantly, *APOBEC1* expression was undetectable in feline liver, lung and MLN RNA-seq samples, and in humans its expression is restricted to the gut (74). If APOBEC1 expression is similar in the cat, it is plausible that intestinal expression of APOBEC1 could introduce C>T mutations during the enteric phase of FCoV infection and thereby drive FCoV evolution towards more efficient replication in monocytes/macrophages, providing the first evidence of host innate immune processes influencing FCoV evolution. Targeted studies of APOBEC- and ADAR-mediated RNA editing of FCoV in feline cells will determine the true extent of the role of these host proteins in FIP pathogenesis.

Similar to SARS-CoV-2 in humans, the ratio of synonymous to non-synonymous substitutions across the entire genome indicated strong purifying selection in all cats (75), with evidence for more relaxed selection in specific genes, particularly S. Multiple mutations were detected within a ∼45 nt region encoding a loop in the HR1 domain of Spike. This domain undergoes a major conformational change to facilitate fusion with the host cell membrane, and it is possible that iSNVs across tissues in this region are associated with tropism change and / or epitopes important in immune evasion. In support of the latter, a previous study identified linear epitopes in the HR1 domain of FCoV-1 Spike protein containing the residues found to be mutated in the present study, which stimulated increased IFN-γ production in PBMCs from FCoV-infected non-FIP cats compared to specific pathogen free (SPF) controls (76). Similarly, SNVs detected in FCoVs from different cats were particularly concentrated within a predicted N-terminal domain of the M gene, which is expected to extend outwards from the viral membrane, and thus is more likely to be accessible to antibodies. This domain is not conserved in other coronaviruses, such as SARS-CoV-2, and therefore could represent an underappreciated immunogenic site on FCoV virions.

Whilst innate immunity might play a role in the acquisition of in-host viral mutations, a robust T cell response early in FCoV infection is believed to protect cats from developing FIP (77,78). The predominant antigen eliciting immune responses in cats with FIP is the FCoV Spike protein, with several studies reporting T cell responses to epitopes derived from FCoV-1 and FCoV-2 Spike (76,79,80). TCR repertoire sequencing has not been applied to feline infectious disease prior to this study. In COVID-19 in humans, TCR repertoire studies demonstrate a skewed profile of TCR V(D)J recombination indicating T cell exhaustion and broad expansion of diverse clonotypes in severe disease (21). In this study, the TCR repertoire was compared between cats with FIP and non-FIP controls. Expansion of clonotypes was significantly higher in the MLN of FIP cats where FCoV protein was detectable by IHC, reducing the overall diversity of the TCR repertoire in these cats. This is consistent with the function of the MLN as a secondary lymphoid organ coordinating the lymphocyte response to circulating foreign antigens, and with FCoV-specific activation of T cell responses in cats with established FIP. Clustered TCRα CDR3 sequences associated with the FIP group showed limited overlap among cats, and clusters that did overlap tended to be highly expanded in only a single cat, even within the group positive for FCoV, indicating that the T cell response is highly polyclonal and divergent among FIP cats. Conversely, several TCRα clusters associated with the control group were highly shared among control cats and detected in very few FIP cats. It is likely that these represent protective clonotypes that effectively restrict FCoV replication, circulating as a consequence of prior exposure to FCoV and recovery in the control group, and may be a consequence of variation in the major histocompatibility complex (MHC), which in cats is poorly understood. Future studies exploring the dynamics of the TCR repertoire at different stages of FIP, including non-FIP cats with systemic FCoV infection and cats recovered from FIP following antiviral therapy, could help to identify effective TCR clonotypes specific for FCoV antigens.

While a strong, directed T-cell response to FCoV is considered protective against development of FIP, the B-cell response has been considered dysregulated and detrimental (81). Clinically, cats with FIP commonly present with marked polyclonal hypergammaglobulinaemia (16), and enhanced uptake of FCoV by feline macrophages in the presence of FCoV-specific antibodies has been demonstrated *in vitro* (82). Antibody-dependent enhancement (ADE) possibly explains why passively immunised cats have an increased likelihood of developing the disease (17,20). Nevertheless, most seropositive cats do not develop FIP, and the relevance of ADE to FIP pathogenesis outside experimental studies remains uncertain.

Study of the feline BCR repertoire has been hindered by the lack of a publicly available, comprehensive library of feline immunoglobulin heavy chain (IGH) VDJ sequences, despite assembly of feline IGH VDJ sequences in a small number of healthy cats (83,84). This is largely due to the ambiguity of this complex region in the most widely used feline genome assembly, felCat9.0. To this end, in the present study, feline IGH VDJ sequences were identified using a combination of *de novo* prediction and manual annotation of the predicted IGH locus in the more recent feline genome assembly fca126, as well as publicly available feline IGH sequences and expression data from the MLN RNA-seq generated in this study.

IGH clonotypes assembled using our newly created feline IGH VDJ reference were more highly expanded in FIP samples compared to controls, and the degree of expansion was highly correlated with FCoV abundance in the MLN, but not the liver or lung. This is likely due to the concentration of germinal centres in the MLN facilitating B cell expansion and diversification. FIP cats differed significantly from control cats in their IGHV gene and isotype usage, suggesting FCoV-dependent changes in VDJ and class-switch recombination. Usage of the IGHG constant gene was higher in all tissues from FIP cats, consistent with the clinical presentation of hypergammaglobulinemia in cats with FIP. Lower somatic hypermutation frequency was observed in IGH clonotypes from FIP cats, which was similar to patterns observed in BCR repertoires from human patients with COVID-19 (22). Concentration of these mutations within complementarity-determining regions (CDRs) also suggested FCoV-driven changes in the feline BCR repertoire through antigen-specific selection.

Inclusion of a cat with systemic coronavirus infection but without FIP (control 164) also provided an opportunity to examine the immune repertoire in this circumstance, and further studies of such cats will be valuable. The IGH repertoire of FCoV-positive control cat 164 was an outlier to the control group in the analysis and appeared more similar to IGH repertoires from FIP cats than controls, suggesting that many of the differences between FIP and control cats are a result of exposure to FCoV, and not necessarily predictive of FIP. It should be noted that the immune repertoire changes with age (85), and in this study, the FIP cats were typically younger than the controls, so an age-related impact could not be ruled out. Nonetheless, the limited degree of between-cat repertoire overlap, in contrast with the high degree of within-cat repertoire overlap, indicates that convergent B-cell responses are rare, and that there is no single immunodominant epitope against which strongly neutralising antibodies are raised in cats with FIP. These findings therefore support the hypothesis that activation of the humoral immune response does not confer protection against FIP, and it remains possible that FCoV-specific antibodies exacerbate disease in the presence of high levels of virus replication through ADE.

The primary limitations of this study are the small cohort size for RNA-seq, and restriction to DSH cats from the UK. In principle, each cat in the FIP group might have been at a different stage of the disease and information on duration of clinical signs was not available for all cats. However, with the widespread adoption of antivirals in the treatment of FIP, availability of such samples will diminish further, and the results of this study provide clear justification for the application of single-cell approaches to tissue or non-invasive samples from FIP cats (e.g. blood) to clarify the role of specific T- and B-cell subsets. Finally, the lack of data on feline BCR/TCR CDR3 epitope specificity precludes prediction of FCoV antigens recognised by the FIP-associated clonotypes identified in this study, but these results provide a solid foundation for the design of future targeted epitope screening experiments relevant to FIP.

Taken together, the results of this study highlight an important role for the host immune response in FIP pathogenesis and reveal for the first time the dynamics and composition of the feline immune repertoire in response to FCoV. Persistent replication of FCoV in spite of pronounced immune activation is a hallmark of FIP, helping to explain why anti-viral therapies work so effectively. However, such therapies carry a risk of inducing anti-viral resistance, so it would be preferable instead to promote an effective immune response. Evaluation of the ineffective adaptive immune response in FIP has revealed remarkably little overlap in dysfunctional TCR and BCR clonotypes. In contrast, the presence of shared TCR clonotypes in non-FIP cats has potential implications for novel therapies, including vaccine design. Further work to characterise the immune repertoire associated with spontaneous recovery from systemic coronavirus infection, or recovery from FIP following anti-viral treatment, could enable identification of at-risk cats, as well as the design of novel targeted immunological interventions for FIP.

## Materials and methods

### Sample collection

Tissue samples (liver, lung, and mesenteric lymph node) were collected post-mortem from cats with or without FIP euthanised for clinical reasons. Full informed consent was obtained from owners, and collection, storage and use of samples were approved by the University of Bristol Animal Welfare and Ethical Review Board (references VIN/14/013, VIN/18/003, VIN/25/005). The study was also approved by the Royal Veterinary College Clinical Research Ethical Review Board (reference URN 2020 1997-2). Samples were archived in RNAlater (Ambion, Inc.) and stored at -80°C for up to 10 years prior to commencement of this study. FIP cases and non-FIP controls were identified based on clinical signs, histological examination and the detection of FCoV antigen within lesional macrophages by immunohistochemistry. Samples were selected based on confidence of diagnosis, and tissues from two cats (142 and 192) with a restriction of FIP lesions to the kidneys were included for comparison. These cats demonstrated acute (cat 142) or chronic (cat 192) FIP lesions in the kidney. At post-mortem examination cat 192 was also found to have chronic inflammation with focal granulomatous infiltrate containing FCoV positive macrophages in the brain. Signalment and diagnostic information for the cats included in the RNA-seq experiment are available in Supplementary Table S1.

### RNA extraction, library preparation and sequencing

For extraction of total RNA, 20-30 mg of feline tissue was added to 600 μl buffer RLT (Qiagen) with 10 μl β-mercaptoethanol in Lysing Matrix D tubes (MP Biomedicals) and homogenised using a FastPrep-24 Classic bead beating grinder and lysis system (MP Biomedicals). RNA was purified from the resulting homogenate using a RNeasy Plus Mini kit (Qiagen), including column-based elimination of genomic DNA, according to the manufacturer’s instructions. RNA purity was assessed using a Nanodrop ND-1000 instrument (ThermoFisher), and RNA integrity was determined using a Tapestation 4200 (Agilent). RNA-seq libraries were prepared from total RNA by Oxford Genomics Centre (Wellcome Centre for Human Genetics, University of Oxford). Libraries were prepared following ribodepletion with the NEBNext rRNA Depletion Kit v2 (NEB) to remove abundant rRNA species. Stranded libraries were generated using a TruSeq Stranded Total RNA kit (Illumina) and sequenced as 150bp paired-end reads on a NovaSeq6000 instrument (Illumina) with a target depth of ∼100M reads per sample.

### Host transcriptome analysis

Raw RNA-seq read quality was assessed using FastQC (86). Adapters were trimmed from raw reads using PEAT (87). Trimmed reads were aligned to the felCat9 reference genome (GCA_000181355.4) using STAR (88), and resulting alignments were filtered with Samtools (89) to retain only properly paired, uniquely mapped reads. Gene-level counts were obtained using the featureCounts function from the Rsubread package (90) with felCat9 gene annotations from Ensembl (91). Counts were normalised using the trimmed mean of M-values (TMM) method in edgeR (92,93) and filtered to remove genes with low expression. Remaining genes were tested for differential expression between FIP and control samples using edgeR, with inclusion of a coefficient for the focal renal FIP samples (cats 142 and 192) in the generalised linear model. Genes were considered differentially expressed if the (Benjamini-Hochberg) adjusted p-value was < 0.05. Principal component analysis was performed on normalised counts using the prcomp function in R. Gene expression was plotted as log_2_ TMM-normalised counts per million mapped reads (CPM). GO enrichment analysis was performed using clusterProfiler (94) with annotated feline gene sets available in the msigdbr package (95) in R/Bioconductor. GO Biological Process gene sets with an FDR-adjusted p-value < 0.05 were considered significantly enriched. For GSEA, feline Hallmark gene sets, also retrieved using the msigdbr package in R, were used with TMM-normalised RNA-seq counts from edgeR in the desktop GSEA application (v4.3.2) from the Broad institute (42,96). Hallmark gene sets with FDR-adjusted p-value < 0.05 were considered significant. For detection of FCoV reads in feline tissue RNA-seq samples, reads were realigned to a hybrid assembly including felCat9 and the full genome of FCoV strain C1Je (DQ848678) using STAR. Counts of reads mapping to the C1Je genome were obtained using featureCounts and plotted as log_2_ TMM-normalised CPM.

### FCoV genome assembly and annotation

Following alignment to the feline reference genome, unmapped reads were extracted from BAM alignments using samtools and subjected to metagenomic assembly using coronaSPAdes (97) with default settings. Single contigs ∼29 kbp with a positive BLAST hit for FCoV were considered candidate full-length FCoV assemblies, and shorter sequences with similarity to public FCoV sequences were considered candidate partial FCoV assemblies. Completeness and contamination of assembled FCoV sequences was assessed using CheckV (98). A summary of the results is available in Supplementary Table S5. Open reading frames (ORFs) in assembled FCoV sequences were detected using Prodigal (99), and annotated according to expected position and sequence similarity to FCoV genes by BLAST search. Where Prodigal failed to annotate overlapping ORFs, as was the case for the E gene in several assemblies, sequences were manually checked for ORFs of similar length and sequence to FCoV E, with a correct start (ATG) and stop codon. For comparisons of sequence identity and phylogenetics, full-length FCoV assemblies were aligned using MUSCLE (v3.8.31) (100). A maximum likelihood tree based on whole genome alignment of candidate FCoV assemblies and FCoV whole genomes retrieved from the NCBI Virus database was produced using RAxML (v8.2.12) (101) with 100 bootstrap replicates.

### Analysis of FCoV intrahost single nucleotide variants (iSNVs)

For detection of iSNVs, the input reads for each assembly were aligned to the FCoV genome assembled from the MLN sample for the same cat, where available, using bwa-mem (102). Variant positions in FCoV reads from each tissue relative to the MLN FCoV assembly for each cat were called using LoFreq (103) with default settings, following insertion of indel scores using lofreq indelqual –dindel. Called single nucleotide variants (SNVs) and indels were annotated using SnpEff (104). Annotated variants were filtered to remove those in the 5’ and 3’ UTRS, and also variants with < 5 reads containing the alternate allele and an alternate allele frequency (AAF) < 0.02. Estimation of dN/dS ratios was performed using the dndscv package (105) in R. For comparison of non-synonymous variants in the FCoV S and M genes across FIP cats, positions of mutated amino acid residues were corrected based on codon-aware alignment of CDS sequences from each cat’s MLN FCoV assembly using codon-msa from the HyPhy package (106). Coordinates of domains in the FCoV Spike protein were determined based on alignment with the FCoV UU4 cryo-EM structure (PDB: 6JX7) published previously (107). Visualisations of FCoV Spike structural models were produced for 6JX7 using ChimeraX (v1.9) (108).

### Feline TCR library preparation and sequencing

For targeted sequencing of the feline TCR repertoire, an expanded cohort of MLN samples was selected from the same feline tissue archive, which included the samples used for the RNA-seq experiment (21 FIP, 24 control). Signalment and TCR sequencing statistics for the cats included in the TCR sequencing experiment are available in Supplementary Table S7. Total RNA was extracted from each MLN sample using the same protocol as for the RNA-seq samples. Due to low A260/230 ratios in some cases, all samples were then cleaned up using an RNA Clean & Concentrator-25 kit (Zymo). Feline TCR libraries were prepared from total RNA as in (23), with modifications. All primer sequences and concentrations used are available in Supplementary Table S8. For each RNA sample, 1 μg total RNA was converted to cDNA by 5’ RACE using SMARTScribe Reverse Transcriptase (Takara Bio) according to the manufacturer’s instructions. Each reaction included 0.5 μM anchored oligo(dT) VN and 1 μΜ template switch oligonucleotide (TSO). Resulting cDNA was cleaned up using AMPure XP beads (Beckman) according to the manufacturer’s instructions and eluted in 40 μl ddH_2_O. Each TCR locus (TRA, TRB, TRG and TRD) was amplified in a separate PCR reaction using KAPA HiFi HotStart ReadyMix (Roche) with the following conditions: 95°C for 3 min, 25 cycles of 98°C for 20 s, 64°C for 15 s, 72°C for 30 s, and a final extension step at 72°C for 3 min. First round PCR products were pooled and purified using AMPure XP beads at 0.8X sample volume and eluted in 50 μl ddH_2_O. 5 μl of the purified product was used in a second round of PCR to incorporate adapter sequences and combinatorial dual indexes according to the instructions available at https://support.illumina.com/content/dam/illumina-support/documents/documentation/chemistry_documentation/16s/16s-metagenomic-library-prep-guide-15044223-b.pdf. Libraries were purified using AMPure XP beads at 0.5X sample volume, eluted in 35 μl ddH_2_O, and library concentrations were determined using a NEBNext Library Quant Kit for Illumina (NEB). Libraries were pooled in equimolar amounts and gel extracted using a QIAquick Gel Extraction Kit (Qiagen) to select fragments at the expected size of ∼600 bp, followed by a final clean up with a MinElute PCR Purification Kit (Qiagen). Feline TCR libraries were sequenced as 300 bp paired-end reads on a MiSeq instrument (Illumina).

### Feline TCR repertoire analysis

Feline TCRα and TCRβ sequences were retrieved from the IMGT database (51) and used to build a reference V(D)J sequence library using the repseqio function in MiXCR (v4.7.0) (109). RNA-seq reads from each sample were assembled into TCRα and TCRβ clonotypes using MiXCR with the rna-seq preset. Resulting TCR clonotypes were analysed with the Immunarch package (110) in R. For targeted TCR sequencing libraries, TCRα and TCRβ clonotypes were assembled using MiXCR with the generic-amplicon preset and the following options: --rna -- rigid-left-alignment-boundary --floating-right-alignment-boundary C --assemble-contigs-by CDR3. Resulting clonotypes were analysed with Immunarch, and clonotypes with CDR3 sequences longer than 30 amino acids were filtered out, as well as samples with a TCR read count < 5,000 (control - 69, 115, FIP - 83, 176). For calculation of TCR repertoire diversity metrics, repertoires were downsampled to the minimum number of reads across samples (TCRα - 7,618, TCRβ - 10,539). Clustering and network analysis of TCR clonotypes was conducted using the NAIR package (111) in R, with TCRs clustered by a maximum Hamming distance of 1 in CDR3 amino acid sequences. Using the findAssociatedSeqs function from NAIR to compare repertoires from FIP and control cats, CDR3 sequences with a p-value < 0.05 (Fisher’s exact test) were considered FIP-associated. Large repertoire networks (>1,000 nodes) were visualised using Gephi (v0.10.1) (112), and individual cluster networks were visualised using igraph (113) and ggraph (114) in R. Generation probability (pGen) for TCR clonotypes was calculated using OLGA (115), with a generative V(D)J recombination model inferred with IGoR (116) based on all non-productive TCR clonotypes from control cats. Sequence logos for FIP-associated TCR clusters were plotted using the ggseqlogo package (117) in R.

### Mapping the feline immunoglobulin heavy chain (IGH) locus

De novo prediction of feline IGH VDJ gene segments was carried out using IgDetective (118) and StochHMM (119) (with models previously used to characterise the canine IGH locus (120)) on the fca126 genome assembly (GCF_018350175.1), which identified 58 IGHV genes (41 functional, 17 non-functional), 7 IGHD genes and 6 IGHJ genes on chromosome B3. A further 7 IGHV genes (5 functional, 2 non-functional) were identified using blastn (121) to identify regions of fca126 chromosome B3 with high sequence identity to the IGHV genes detected using both IgDetective and StochHMM. All genes were manually checked for intact and correctly spaced 7mer/9mer motifs of the recombination signal sequence, as well as expression in feline MLN RNA-seq samples using the Integrative Genomics Viewer (IGV) (122). Constant region genes were identified by using blastn to detect constant region sequences from previously sequenced feline IGH clonotypes (83,123), as well as canine IGH constant region genes retrieved from IMGT, in fca126. Multiple sequence alignment of the 65 IGHV genes identified in fca126 revealed 4 subgroups of IGHV genes, 3 of which had high sequence identity (≥ 75%) with canine IGHV gene subgroups. The IGHV subgroup with no significant sequence identity to any canine IGHV genes, comprising a single pseudogene, was tentatively named IGHV(I)-22, while all other detected feline IGHV genes were named according to the IMGT nomenclature rules (124). Genomic coordinates of all identified IGH VDJ genes relative to fca126 were determined using blastn. For downstream analysis, a reference library in JSON format was created using candidate VDJ and constant region gene sequences in FASTA format with the repseqio function in MiXCR, and all sequences used are available in Supplementary Table S9.

### Feline BCR repertoire analysis

RNA-seq reads from each feline tissue sample were assembled into IGH clonotypes using MiXCR with the rna-seq preset and the IGH VDJ sequence reference library described in the previous section. Resulting IGH clonotypes were analysed with Immunarch in R. Clonotypes with CDR3 sequences longer than 35 amino acids were filtered out, as well as samples with low IGH read counts (…). For calculation of IGH repertoire diversity metrics, repertoires were downsampled to the minimum number of reads across samples (51,568). For visualisation of representative BCR repertoire networks for each sample, a network of all IGH clonotypes was constructed using the NAIR package in R, with a maximum Hamming distance of 1 in CDR3 nucleotide sequences. Each network was then downsampled to 2,000 clusters 1,000 times, and one subsample with a maximum cluster size closest to the median of all subsamples was randomly selected and visualised using igraph and ggraph. Repertoire overlap was determined by constructing a global network with all IGH clonotypes assembled for all samples, and a maximum Hamming distance of 1 in CDR3 amino acid sequences. Degree of sample overlap for each cluster was then determined by the Jaccard index. Networks for IGH clusters detected in multiple samples were visualised using igraph and ggraph, and sequence logos were plotted using the ggseqlogo package in R. For analysis of somatic hypermutation, full-length IGH clonotypes were assembled using MiXCR with the option --assemble-contigs-by VDJ. The effect of allelic variation was accounted for using mixcr findAlleles, and clonotypes with VDJ genes reassigned to identified alleles were converted to AIRR format using mixcr exportAirr. Analysis of mutations in reassigned IGH clonotypes relative to germline sequence was conducted using the shazam R package (125) from the Immcantation framework. Posterior probability density functions and associated selection scores for IGH V regions were calculated based on observed and expected mutation frequencies with the calcBaseline function in shazam.

## Acknowledgements

The authors thank all the practitioners, owners, and colleagues who helped in the acquisition of samples used in this study. The authors also thank Dr Rachael Bashford-Rogers and Dr Lauren Overend for helpful discussions concerning TCR/BCR sequencing and analysis, and Dr Angela Lee and the staff at Oxford Genomics Centre for their support. Computation used the Oxford Biomedical Research Computing (BMRC) facility, a joint development between the Wellcome Centre for Human Genetics and the Big Data Institute supported by Health Data Research UK and the NIHR Oxford Biomedical Research Centre. The views expressed are those of the author(s) and not necessarily those of the NHS, the NIHR or the Department of Health.

## Author contributions

Conceptualization - L.J.D., T.K.H., MASCOT

Data curation - T.K.H., E.N.B.

Formal Analysis - T.K.H.

Funding acquisition - L.J.D., T.K.H., MASCOT

Investigation - T.K.H., S.F., A.K.

Methodology - T.K.H.

Project administration - L.J.D., T.K.H., S.F., E.N.B., M.J.H.

Resources - L.J.D., E.N.B., M.J.H., C.A.O.

Software - T.K.H. Supervision - L.J.D., C.A.O.

Validation - L.J.D., T.K.H. Visualization - T.K.H.

Writing (original draft) - T.K.H., L.J.D.

Writing (review & editing) - T.K.H., L.J.D., MASCOT, A.K., M.J.H., C.A.O., E.N.B.

## Data availability

Sequencing data supporting the findings of this study have been deposited in the National Center for Biotechnology Information (NCBI) Sequence Read Archive under BioProject ID PRJNA1330231. Full-length FCoV genome assemblies have been deposited in NCBI GenBank under accession numbers PX315748-PX315758 inclusive. All other data needed to evaluate the conclusions in the paper are present in the paper and/or the Supplementary Materials.

## The MASCOT Consortium

In addition to the authors, consortium members are listed in alphabetical order.

Sophie Binks^1,2^, Katherine Hughes^3^, Lorna J. Kennedy^4^, Emily Kwan^5^, Judy A. Mitchell^6^, Androniki Psifidi^5^, Rachael E. Tarlinton^7^, Marsha D. Wallace^5,8^

^1^ Oxford Autoimmune Neurology Group, Nuffield Department of Clinical Neuroscience, Oxford, UK

^2^ Department of Neurology, John Radcliffe Hospital, Oxford, UK

^3^ Department of Veterinary Medicine, University of Cambridge, Cambridge, UK

^4^ Centre for Integrated Genomic Medical Research, University of Manchester, Manchester, UK

^5^ Department of Clinical Science and Services, The Royal Veterinary College, Hatfield, UK

^6^ Department of Pathobiology & Population Sciences, The Royal Veterinary College, Hatfield, UK

^7^ School of Veterinary Medicine and Science, University of Nottingham, Loughborough, UK

^8^ Department of Physiology, Anatomy and Genetics, University of Oxford, OX1 3PT, UK

## References

1. Kipar A, May H, Menger S, Weber M, Leukert W, Reinacher M. Morphologic Features and Development of Granulomatous Vasculitis in Feline Infectious Peritonitis. Vet Pathol. 2005 May 1;42(3):321–30.

2. Kipar A, Meli ML. Feline Infectious Peritonitis: Still an Enigma? Vet Pathol. 2014 Mar 1;51(2):505–26.

3. Taylor SS, Coggins S, Barker EN, Gunn-Moore D, Jeevaratnam K, Norris JM, et al. Retrospective study and outcome of 307 cats with feline infectious peritonitis treated with legally sourced veterinary compounded preparations of remdesivir and GS-441524 (2020–2022). J Feline Med Surg. 2023 Sep 1;25(9):1098612X231194460.

4. Murphy BG, Perron M, Murakami E, Bauer K, Park Y, Eckstrand C, et al. The nucleoside analog GS-441524 strongly inhibits feline infectious peritonitis (FIP) virus in tissue culture and experimental cat infection studies. Vet Microbiol. 2018 Jun 1;219:226–33.

5. Zuzzi-Krebitz AM, Buchta K, Bergmann M, Krentz D, Zwicklbauer K, Dorsch R, et al. Short Treatment of 42 Days with Oral GS-441524 Results in Equal Efficacy as the Recommended 84-Day Treatment in Cats Suffering from Feline Infectious Peritonitis with Effusion—A Prospective Randomized Controlled Study. Viruses. 2024 Jul;16(7):1144.

6. Roy M, Jacque N, Novicoff W, Li E, Negash R, Evans SJM. Unlicensed Molnupiravir is an Effective Rescue Treatment Following Failure of Unlicensed GS-441524-like Therapy for Cats with Suspected Feline Infectious Peritonitis. Pathogens. 2022 Oct;11(10):1209.

7. Renner KA, Cattin R, Kimble B, Munday J, White A, Coggins S. Efficacy of oral remdesivir in treating feline infectious peritonitis: a prospective observational study of 29 cats. J Feline Med Surg. 2025 May 1;27(5):1098612X251335189.

8. Tasker S, Addie DD, Egberink H, Hofmann-Lehmann R, Hosie MJ, Truyen U, et al. Feline Infectious Peritonitis: European Advisory Board on Cat Diseases Guidelines. Viruses. 2023 Sep;15(9):1847.

9. Worthing KA, Wigney DI, Dhand NK, Fawcett A, McDonagh P, Malik R, et al. Risk factors for feline infectious peritonitis in Australian cats. J Feline Med Surg. 2012 Jun 1;14(6):405–12.

10. Pesteanu-Somogyi LD, Radzai C, Pressler BM. Prevalence of feline infectious peritonitis in specific cat breeds★. J Feline Med Surg. 2006 Feb 1;8(1):1–5.

11. Golovko L, Lyons LA, Liu H, Sørensen A, Wehnert S, Pedersen NC. Genetic susceptibility to feline infectious peritonitis in Birman cats. Virus Res. 2013 Jul 1;175(1):58–63.

12. Malbon AJ, Russo G, Burgener C, Barker EN, Meli ML, Tasker S, et al. The Effect of Natural Feline Coronavirus Infection on the Host Immune Response: A Whole-Transcriptome Analysis of the Mesenteric Lymph Nodes in Cats with and without Feline Infectious Peritonitis. Pathogens. 2020 Jul;9(7):524.

13. Garner MM, Ramsell K, Morera N, Juan-Sallés C, Jiménez J, Ardiaca M, et al. Clinicopathologic Features of a Systemic Coronavirus-Associated Disease Resembling Feline Infectious Peritonitis in the Domestic Ferret (Mustela putorius). Vet Pathol. 2008 Mar 1;45(2):236–46.

14. Decaro N, Martella V, Elia G, Campolo M, Desario C, Cirone F, et al. Molecular characterisation of the virulent canine coronavirus CB/05 strain. Virus Res. 2007 Apr 1;125(1):54–60.

15. Sweet AN, André NM, Stout AE, Licitra BN, Whittaker GR. Clinical and Molecular Relationships between COVID-19 and Feline Infectious Peritonitis (FIP). Viruses. 2022 Mar;14(3):481.

16. Paltrinieri S, Cammarata MP, Cammarata G, Comazzi S. Some aspects of humoral and cellular immunity in naturally occuring feline infectious peritonitis. Vet Immunol Immunopathol. 1998 Oct 23;65(2):205–20.

17. Takano T, Yamada S, Doki T, Hohdatsu T. Pathogenesis of oral type I feline infectious peritonitis virus (FIPV) infection: Antibody-dependent enhancement infection of cats with type I FIPV via the oral route. J Vet Med Sci. 2019;81(6):911–5.

18. Poland AM, Vennema H, Foley JE, Pedersen NC. Two related strains of feline infectious peritonitis virus isolated from immunocompromised cats infected with a feline enteric coronavirus. J Clin Microbiol. 1996 Dec;34(12):3180–4.

19. Pedersen NC. A review of feline infectious peritonitis virus infection: 1963–2008. J Feline Med Surg. 2009 Apr 1;11(4):225–58.

20. Takano T, Kawakami C, Yamada S, Satoh R, Hohdatsu T. Antibody-Dependent Enhancement Occurs Upon Re-Infection with the Identical Serotype Virus in Feline Infectious Peritonitis Virus Infection. J Vet Med Sci. 2008;70(12):1315–21.

21. Park JJ, Lee KAV, Lam SZ, Moon KS, Fang Z, Chen S. Machine learning identifies T cell receptor repertoire signatures associated with COVID-19 severity. Commun Biol. 2023 Jan 20;6(1):76.

22. Galson JD, Schaetzle S, Bashford-Rogers RJM, Raybould MIJ, Kovaltsuk A, Kilpatrick GJ, et al. Deep Sequencing of B Cell Receptor Repertoires From COVID-19 Patients Reveals Strong Convergent Immune Signatures. Front Immunol [Internet]. 2020 Dec 15 [cited 2025 Jul 11];11. Available from: https://www.frontiersin.org/journals/immunology/articles/10.3389/fimmu.2020.605170/full

23. Radtanakatikanon A, Keller SM, Darzentas N, Moore PF, Folch G, Nguefack Ngoune V, et al. Topology and expressed repertoire of the Felis catus T cell receptor loci. BMC Genomics. 2020 Jan 6;21(1):20.

24. Gao YY, Wang Q, Liang XY, Zhang S, Bao D, Zhao H, et al. An updated review of feline coronavirus: mind the two biotypes. Virus Res. 2023 Mar 1;326:199059.

25. 25. Intra-host variation in the spike S1/S2 region of a feline coronavirus type-1 in a cat with persistent infection | bioRxiv [Internet]. [cited 2025 Jul 15]. Available from: https://www.biorxiv.org/content/10.1101/2023.07.31.551356v1

26. Chang HW, Egberink HF, Rottier PJM. Sequence Analysis of Feline Coronaviruses and the Circulating Virulent/Avirulent Theory. Emerg Infect Dis. 2011 Apr;17(4):744–6.

27. Brown MA, Troyer JL, Pecon-Slattery J, Roelke ME, O’Brien SJ. Genetics and Pathogenesis of Feline Infectious Peritonitis Virus - Volume 15, Number 9—September 2009 - Emerging Infectious Diseases journal - CDC. [cited 2025 Jul 15]; Available from: https://wwwnc.cdc.gov/eid/article/15/9/08-1573_article

28. Attipa C, Warr AS, Epaminondas D, O’Shea M, Hanton AJ, Fletcher S, et al. Feline infectious peritonitis epizootic caused by a recombinant coronavirus. Nature. 2025 Jul 9;1–3.

29. Herrewegh AAPM, Smeenk I, Horzinek MC, Rottier PJM, de Groot RJ. Feline Coronavirus Type II Strains 79-1683 and 79-1146 Originate from a Double Recombination between Feline Coronavirus Type I and Canine Coronavirus. J Virol. 1998 May;72(5):4508–14.

30. Lednicky JA, Tagliamonte MS, White SK, Blohm GM, Alam MM, Iovine NM, et al. Isolation of a Novel Recombinant Canine Coronavirus From a Visitor to Haiti: Further Evidence of Transmission of Coronaviruses of Zoonotic Origin to Humans. Clin Infect Dis. 2022 Jul 1;75(1):e1184–7.

31. Vlasova AN, Diaz A, Damtie D, Xiu L, Toh TH, Lee JSY, et al. Novel Canine Coronavirus Isolated from a Hospitalized Patient With Pneumonia in East Malaysia. Clin Infect Dis. 2022 Jan 1;74(3):446–54.

32. Barker EN, Stranieri A, Helps CR, Porter EL, Davidson AD, Day MJ, et al. Limitations of using feline coronavirus spike protein gene mutations to diagnose feline infectious peritonitis. Vet Res. 2017 Oct 5;48(1):60.

33. Li C, Liu Q, Kong F, Guo D, Zhai J, Su M, et al. Circulation and genetic diversity of Feline coronavirus type I and II from clinically healthy and FIP-suspected cats in China. Transbound Emerg Dis. 2019 Mar 1;66(2):763–75.

34. Kummrow M, Meli ML, Haessig M, Goenczi E, Poland A, Pedersen NC, et al. Feline Coronavirus Serotypes 1 and 2: Seroprevalence and Association with Disease in Switzerland. Clin Vaccine Immunol [Internet]. 2005 Oct [cited 2025 Sep 30]; Available from: https://journals.asm.org/doi/10.1128/CDLI.12.10.1209-1215.2005

35. Benetka V, Kübber-Heiss A, Kolodziejek J, Nowotny N, Hofmann-Parisot M, Möstl K. Prevalence of feline coronavirus types I and II in cats with histopathologically verified feline infectious peritonitis. Vet Microbiol. 2004 Mar 26;99(1):31–42.

36. Chang HW, Egberink HF, Halpin R, Spiro DJ, Rottier PJM. Spike Protein Fusion Peptide and Feline Coronavirus Virulence - Volume 18, Number 7—July 2012 - Emerging Infectious Diseases journal - CDC. [cited 2025 Jul 10]; Available from: https://wwwnc.cdc.gov/eid/article/18/7/12-0143_article

37. Licitra BN, Millet JK, Regan AD, Hamilton BS, Rinaldi VD, Duhamel GE, et al. Mutation in Spike Protein Cleavage Site and Pathogenesis of Feline Coronavirus - Volume 19, Number 7—July 2013 - Emerging Infectious Diseases journal - CDC. [cited 2025 Jul 10]; Available from: https://wwwnc.cdc.gov/eid/article/19/7/12-1094_article

38. Thayer V, Gogolski S, Felten S, Hartmann K, Kennedy M, Olah GA. 2022 AAFP/EveryCat Feline Infectious Peritonitis Diagnosis Guidelines. J Feline Med Surg. 2022 Sep 1;24(9):905–33.

39. Slaviero M, Cony FG, da Silva RC, De Lorenzo C, de Almeida BA, Bertolini M, et al. Pathological findings and patterns of feline infectious peritonitis in the respiratory tract of cats. J Comp Pathol. 2024 Apr 1;210:15–24.

40. Samarajiwa SA, Forster S, Auchettl K, Hertzog PJ. INTERFEROME: the database of interferon regulated genes. Nucleic Acids Res. 2009 Jan 1;37(suppl_1):D852–7.

41. Liberzon A, Subramanian A, Pinchback R, Thorvaldsdóttir H, Tamayo P, Mesirov JP. Molecular signatures database (MSigDB) 3.0. Bioinformatics. 2011 Jun 15;27(12):1739–40.

42. Subramanian A, Tamayo P, Mootha VK, Mukherjee S, Ebert BL, Gillette MA, et al. Gene set enrichment analysis: A knowledge-based approach for interpreting genome-wide expression profiles. Proc Natl Acad Sci. 2005 Oct 25;102(43):15545–50.

43. Liberzon A, Birger C, Thorvaldsdóttir H, Ghandi M, Mesirov JP, Tamayo P. The Molecular Signatures Database Hallmark Gene Set Collection. Cell Syst. 2015 Dec 23;1(6):417– 25.

44. Malbon AJ, Fonfara S, Meli ML, Hahn S, Egberink H, Kipar A. Feline Infectious Peritonitis as a Systemic Inflammatory Disease: Contribution of Liver and Heart to the Pathogenesis. Viruses. 2019 Dec;11(12):1144.

45. Transcriptional profiling of feline infectious peritonitis virus infection in CRFK cells and in PBMCs from FIP diagnosed cats | Virology Journal | Full Text [Internet]. [cited 2025 Jul 10]. Available from: https://virologyj.biomedcentral.com/articles/10.1186/1743-422X-10-329

46. Dye C, Siddell SG. Genomic RNA sequence of feline coronavirus strain FCoV C1Je. J Feline Med Surg. 2007 Jun 1;9(3):202–13.

47. Phylogenetic Analysis of Feline Coronavirus Strains in an Epizootic Outbreak of Feline Infectious Peritonitis. [cited 2025 Jul 15]; Available from: https://onlinelibrary.wiley.com/doi/10.1111/jvim.12058

48. Malbon AJ, Meli ML, Barker EN, Davidson AD, Tasker S, Kipar A. Inflammatory Mediators in the Mesenteric Lymph Nodes, Site of a Possible Intermediate Phase in the Immune Response to Feline Coronavirus and the Pathogenesis of Feline Infectious Peritonitis? J Comp Pathol. 2019 Jan 1;166:69–86.

49. Glanville J, Huang H, Nau A, Hatton O, Wagar LE, Rubelt F, et al. Identifying specificity groups in the T cell receptor repertoire. Nature. 2017 Jul;547(7661):94–8.

50. Dash P, Fiore-Gartland AJ, Hertz T, Wang GC, Sharma S, Souquette A, et al. Quantifiable predictive features define epitope-specific T cell receptor repertoires. Nature. 2017 Jul;547(7661):89–93.

51. Manso T, Folch G, Giudicelli V, Jabado-Michaloud J, Kushwaha A, Nguefack Ngoune V, et al. IMGT® databases, related tools and web resources through three main axes of research and development. Nucleic Acids Res. 2022 Jan 7;50(D1):D1262–72.

52. Pan H, Peto R, Restrepo AMH, Preziosi MP, Sathiyamoorthy V, Karim QA, et al. Remdesivir and three other drugs for hospitalised patients with COVID-19: final results of the WHO Solidarity randomised trial and updated meta-analyses. The Lancet. 2022 May 21;399(10339):1941–53.

53. Sariol A, Perlman S. Lessons for COVID-19 Immunity from Other Coronavirus Infections. Immunity. 2020 Aug 18;53(2):248–63.

54. Zhang Q, Bastard P, Liu Z, Le Pen J, Moncada-Velez M, Chen J, et al. Inborn errors of type I IFN immunity in patients with life-threatening COVID-19. Science. 2020 Oct 23;370(6515):eabd4570.

55. Konno Y, Kimura I, Uriu K, Fukushi M, Irie T, Koyanagi Y, et al. SARS-CoV-2 ORF3b Is a Potent Interferon Antagonist Whose Activity Is Increased by a Naturally Occurring Elongation Variant. Cell Rep [Internet]. 2020 Sep 22 [cited 2025 Jul 10];32(12). Available from: https://www.cell.com/cell-reports/abstract/S2211-1247(20)31174-8

56. Li JY, Liao CH, Wang Q, Tan YJ, Luo R, Qiu Y, et al. The ORF6, OR F8 and nucleocapsid proteins of SARS-CoV-2 inhibit type I interferon signaling pathway. Virus Res. 2020 Sep 1;286:198074.

57. Goitsuka R, Ohashi T, Ono K, Yasukawa K, Koishibara Y, Fukui H, et al. IL-6 activity in feline infectious peritonitis. J Immunol. 1990 Apr 1;144(7):2599–603.

58. Lucas C, Wong P, Klein J, Castro TBR, Silva J, Sundaram M, et al. Longitudinal analyses reveal immunological misfiring in severe COVID-19. Nature. 2020 Aug;584(7821):463–9.

59. Takano T, Hohdatsu T, Hashida Y, Kaneko Y, Tanabe M, Koyama H. A “possible” involvement of TNF-alpha in apoptosis induction in peripheral blood lymphocytes of cats with feline infectious peritonitis. Vet Microbiol. 2007 Jan 31;119(2):121–31.

60. Haagmans BL, Egberink HF, Horzinek MC. Apoptosis and T-cell depletion during feline infectious peritonitis. J Virol. 1996 Dec;70(12):8977–83.

61. Kiss I, Poland AM, Pedersen NC. Disease outcome and cytokine responses in cats immunized with an avirulent feline infectious peritonitis virus (FIPV)-UCD1 and challenge-exposed with virulent FIPV-UCD8. J Feline Med Surg. 2004 Apr 1;6(2):89– 97.

62. Berg AL, Ekman K, Belák S, Berg M. Cellular composition and interferon-γ expression of the local inflammatory response in feline infectious peritonitis (FIP). Vet Microbiol. 2005 Nov 30;111(1):15–23.

63. Gelain ME, Meli M, Paltrinieri S. Whole blood cytokine profiles in cats infected by feline coronavirus and healthy non-FCoV infected specific pathogen-free cats. J Feline Med Surg. 2006 Dec 1;8(6):389–99.

64. Huang HC, Wang SH, Fang GC, Chou WC, Liao CC, Sun CP, et al. Upregulation of PD- L1 by SARS-CoV-2 promotes immune evasion. J Med Virol. 2023;95(2):e28478.

65. Sun C, Mezzadra R, Schumacher TN. Regulation and Function of the PD-L1 Checkpoint. Immunity. 2018 Mar 20;48(3):434–52.

66. Wykes MN, Lewin SR. Immune checkpoint blockade in infectious diseases. Nat Rev Immunol. 2018 Feb;18(2):91–104.

67. Development of anti-feline PD-1 antibody and its functional analysis | Scientific Reports [Internet]. [cited 2025 Jul 10]. Available from: https://www.nature.com/articles/s41598-023-31543-6

68. Felten S, Klein-Richers U, Unterer S, Bergmann M, Zablotski Y, Hofmann-Lehmann R, et al. Patterns of Feline Coronavirus Shedding and Associated Factors in Cats from Breeding Catteries. Viruses. 2023 Jun;15(6):1279.

69. Porter E, Tasker S, Day MJ, Harley R, Kipar A, Siddell SG, et al. Amino acid changes in the spike protein of feline coronavirus correlate with systemic spread of virus from the intestine and not with feline infectious peritonitis. Vet Res. 2014 Apr 25;45(1):49.

70. Tortorici MA, Choi A, Gibson CA, Lee J, Brown JT, Stewart C, et al. Loss of FCoV-23 spike domain 0 enhances fusogenicity and entry kinetics. Nature. 2025 Jul 9;1–9.

71. Olarte-Castillo XA, Choi A, Frazier LE, Whittaker G. Rethinking the Drivers of Coronavirus Virulence and Pathogenesis: Toward an Understanding of the Dynamic World of Mutations, Indels, and Recombination Within the Species Alphacoronavirus-1. Qeios [Internet]. 2024 Nov 26 [cited 2025 Jul 10]; Available from: https://www.qeios.com/read/YYO05O

72. Cook S, Castillo D, Williams S, Haake C, Murphy B. Serotype I and II Feline Coronavirus Replication and Gene Expression Patterns of Feline Cells—Building a Better Understanding of Serotype I FIPV Biology. Viruses. 2022 Jul;14(7):1356.

73. Evidence for host-dependent RNA editing in the transcriptome of SARS-CoV-2 | Science Advances [Internet]. [cited 2025 Jul 10]. Available from: https://www.science.org/doi/10.1126/sciadv.abb5813

74. Hadjiagapiou C, Giannoni F, Funahashi T, Skarosi SF, Davidson NO. Molecular cloning of a human small intestinal apolipoprotein B mRNA editing protein. Nucleic Acids Res. 1994 May 25;22(10):1874–9.

75. Patterns of within-host genetic diversity in SARS-CoV-2 | eLife [Internet]. [cited 2025 Jul 11]. Available from: https://elifesciences.org/articles/66857

76. Satoh R, Furukawa T, Kotake M, Takano T, Motokawa K, Gemma T, et al. Screening and identification of T helper 1 and linear immunodominant antibody-binding epitopes in the spike 2 domain and the nucleocapsid protein of feline infectious peritonitis virus. Vaccine. 2011 Feb 17;29(9):1791–800.

77. Natural History of a Recurrent Feline Coronavirus Infection and the Role of Cellular Immunity in Survival and Disease | Journal of Virology [Internet]. [cited 2025 Jul 11]. Available from: https://journals.asm.org/doi/10.1128/jvi.79.2.1036-1044.2005?url_ver=Z39.88-2003&rfr_id=ori%3Arid%3Acrossref.org&rfr_dat=cr_pub++0pubmed

78. Paltrinieri S, Ponti W, Comazzi S, Giordano A, Poli G. Shifts in circulating lymphocyte subsets in cats with feline infectious peritonitis (FIP): pathogenic role and diagnostic relevance. Vet Immunol Immunopathol. 2003 Dec 15;96(3):141–8.

79. Chawla M, Cuspoca AF, Akthar N, Magdaleno JSL, Rattanabunyong S, Suwattanasophon C, et al. Immunoinformatics-aided rational design of a multi-epitope vaccine targeting feline infectious peritonitis virus. Front Vet Sci. 2023 Dec 13;10:1280273.

80. Takano T, Tomiyama Y, Katoh Y, Nakamura M, Satoh R, Hohdatsu T. Mutation of neutralizing/antibody-dependent enhancing epitope on spike protein and 7b gene of feline infectious peritonitis virus: Influences of viral replication in monocytes/macrophages and virulence in cats. Virus Res. 2011 Mar 1;156(1):72–80.

81. A review of feline infectious peritonitis virus infection: 1963–2008 - Niels C. Pedersen, 2009 [Internet]. [cited 2025 Jul 11]. Available from: https://journals.sagepub.com/doi/10.1016/j.jfms.2008.09.008?url_ver=Z39.88-2003&rfr_id=ori:rid:crossref.org&rfr_dat=cr_pub%20%200pubmed

82. Antibody-dependent enhancement of serotype II feline enteric coronavirus infection in primary feline monocytes | Archives of Virology [Internet]. [cited 2025 Jul 11]. Available from: https://link.springer.com/article/10.1007/s00705-017-3489-8

83. Lu Z, Tallmadge RL, Callaway HM, Felippe MJB, Parker JSL. Sequence analysis of feline immunoglobulin mRNAs and the development of a felinized monoclonal antibody specific to feline panleukopenia virus. Sci Rep. 2017 Oct 5;7(1):12713.

84. Steiniger SCJ, Glanville J, Harris DW, Wilson TL, Ippolito GC, Dunham SA. Comparative analysis of the feline immunoglobulin repertoire. Biologicals. 2017 Mar 1;46:81–7.

85. Hu J, Pan M, Reid B, Tworoger S, Li B. Quantifiable blood TCR repertoire components associate with immune aging. Nat Commun. 2024 Sep 17;15(1):8171.

86. Andrews S. FastQC: A Quality Control tool for High Throughput Sequence Data [Internet]. 2010 [cited 2025 Jun 23]. Available from: https://www.bioinformatics.babraham.ac.uk/projects/fastqc/

87. Li YL, Weng JC, Hsiao CC, Chou MT, Tseng CW, Hung JH. PEAT: an intelligent and efficient paired-end sequencing adapter trimming algorithm. BMC Bioinformatics. 2015 Dec;16(S1):S2.

88. Dobin A, Davis CA, Schlesinger F, Drenkow J, Zaleski C, Jha S, et al. STAR: ultrafast universal RNA-seq aligner. Bioinformatics. 2013 Jan 1;29(1):15–21.

89. Danecek P, Bonfield JK, Liddle J, Marshall J, Ohan V, Pollard MO, et al. Twelve years of SAMtools and BCFtools. GigaScience. 2021 Jan 29;10(2):giab008.

90. Liao Y, Smyth GK, Shi W. The R package Rsubread is easier, faster, cheaper and better for alignment and quantification of RNA sequencing reads. Nucleic Acids Res. 2019 May 7;47(8):e47–e47.

91. Dyer SC, Austine-Orimoloye O, Azov AG, Barba M, Barnes I, Barrera-Enriquez VP, et al. Ensembl 2025. Nucleic Acids Res. 2025 Jan 6;53(D1):D948–57.

92. Robinson MD, McCarthy DJ, Smyth GK. edgeR : a Bioconductor package for differential expression analysis of digital gene expression data. Bioinformatics. 2010 Jan 1;26(1):139–40.

93. McCarthy DJ, Chen Y, Smyth GK. Differential expression analysis of multifactor RNA-Seq experiments with respect to biological variation. Nucleic Acids Res. 2012 May 1;40(10):4288–97.

94. Wu T, Hu E, Xu S, Chen M, Guo P, Dai Z, et al. clusterProfiler 4.0: A universal enrichment tool for interpreting omics data. The Innovation. 2021 Aug 28;2(3):100141.

95. Dolgalev I. msigdbr: MSigDB Gene Sets for Multiple Organisms in a Tidy Data Format [Internet]. 2025. Available from: https://CRAN.R-project.org/package=msigdbr

96. Mootha VK, Lindgren CM, Eriksson KF, Subramanian A, Sihag S, Lehar J, et al. PGC-1α-responsive genes involved in oxidative phosphorylation are coordinately downregulated in human diabetes. Nat Genet. 2003 Jul;34(3):267–73.

97. Meleshko D, Hajirasouliha I, Korobeynikov A. coronaSPAdes: from biosynthetic gene clusters to RNA viral assemblies. Bioinformatics. 2021 Dec 22;38(1):1–8.

98. Nayfach S, Camargo AP, Schulz F, Eloe-Fadrosh E, Roux S, Kyrpides NC. CheckV assesses the quality and completeness of metagenome-assembled viral genomes. Nat Biotechnol. 2021 May;39(5):578–85.

99. Hyatt D, Chen GL, LoCascio PF, Land ML, Larimer FW, Hauser LJ. Prodigal: prokaryotic gene recognition and translation initiation site identification. BMC Bioinformatics. 2010 Mar 8;11(1):119.

100. Edgar RC. MUSCLE: multiple sequence alignment with high accuracy and high throughput. Nucleic Acids Res. 2004 Mar 1;32(5):1792–7.

101. Stamatakis A. RAxML version 8: a tool for phylogenetic analysis and post-analysis of large phylogenies. Bioinformatics. 2014 May 1;30(9):1312–3.

102. Li H, Durbin R. Fast and accurate short read alignment with Burrows–Wheeler transform. Bioinformatics. 2009 Jul 15;25(14):1754–60.

103. Wilm A, Aw PPK, Bertrand D, Yeo GHT, Ong SH, Wong CH, et al. LoFreq: a sequence-quality aware, ultra-sensitive variant caller for uncovering cell-population heterogeneity from high-throughput sequencing datasets. Nucleic Acids Res. 2012 Dec 1;40(22):11189–201.

104. Cingolani P, Platts A, Wang LL, Coon M, Nguyen T, Wang L, et al. A program for annotating and predicting the effects of single nucleotide polymorphisms, SnpEff: SNPs in the genome of Drosophila melanogaster strain w^1118^ ; iso-2; iso-3. Fly (Austin). 2012 Apr;6(2):80–92.

105. Martincorena I, Raine KM, Gerstung M, Dawson KJ, Haase K, Loo PV, et al. Universal Patterns of Selection in Cancer and Somatic Tissues. Cell. 2017 Nov 16;171(5):1029–1041.e21.

106. Pond SLK, Frost SDW, Muse SV. HyPhy: hypothesis testing using phylogenies. Bioinformatics. 2005 Mar 1;21(5):676–9.

107. Yang TJ, Chang YC, Ko TP, Draczkowski P, Chien YC, Chang YC, et al. Cryo-EM analysis of a feline coronavirus spike protein reveals a unique structure and camouflaging glycans. Proc Natl Acad Sci. 2020 Jan 21;117(3):1438–46.

108. Meng EC, Goddard TD, Pettersen EF, Couch GS, Pearson ZJ, Morris JH, et al. UCSF ChimeraX: Tools for structure building and analysis. Protein Sci. 2023;32(11):e4792.

109. Bolotin DA, Poslavsky S, Mitrophanov I, Shugay M, Mamedov IZ, Putintseva EV, et al. MiXCR: software for comprehensive adaptive immunity profiling. Nat Methods. 2015 May;12(5):380–1.

110. Nazarov VI, Tsvetkov VO, Fiadziushchanka S, Rumynskiy E, Popov AA, Balashov I, et al. immunarch: Bioinformatics Analysis of T-Cell and B-Cell Immune Repertoires. 2023.

111. Yang H, Cham J, Neal BP, Fan Z, He T, Zhang L. NAIR: Network Analysis of Immune Repertoire. Front Immunol [Internet]. 2023 Jul 7 [cited 2025 Jun 23];14. Available from: https://www.frontiersin.org/journals/immunology/articles/10.3389/fimmu.2023.1181825/f ull

112. Mathieu Bastian SH. Gephi: An Open Source Software for Exploring and Manipulating Networks [Internet]. AAAI. [cited 2025 Jun 23]. Available from: https://aaai.org/papers/00361-13937-gephi-an-open-source-software-for-exploring-and-manipulating-networks/

113. Csárdi G, Nepusz T, Traag V, Horvát S, Zanini F, Noom D, et al. igraph: Network Analysis and Visualization in R [Internet]. 2025. Available from: https://CRAN.R-project.org/package=igraph

114. Pedersen TL. ggraph: An Implementation of Grammar of Graphics for Graphs and Networks [Internet]. 2024. Available from: https://ggraph.data-imaginist.com

115. Sethna Z, Elhanati Y, Callan CG Jr, Walczak AM, Mora T. OLGA: fast computation of generation probabilities of B- and T-cell receptor amino acid sequences and motifs. Bioinformatics. 2019 Sep 1;35(17):2974–81.

116. Marcou Q, Mora T, Walczak AM. High-throughput immune repertoire analysis with IGoR. Nat Commun. 2018 Feb 8;9(1):561.

117. Wagih O. ggseqlogo: a versatile R package for drawing sequence logos. Bioinformatics. 2017 Nov 15;33(22):3645–7.

118. Sirupurapu V, Safonova Y, Pevzner PA. Gene prediction in the immunoglobulin loci. Genome Res. 2022 Jan 6;32(6):1152–69.

119. Lott PC, Korf I. StochHMM: a flexible hidden Markov model tool and C++ library. Bioinformatics. 2014 Jun 1;30(11):1625–6.

120. Hwang MH, Darzentas N, Bienzle D, Moore PF, Morrison J, Keller SM. Characterization of the canine immunoglobulin heavy chain repertoire by next generation sequencing. Vet Immunol Immunopathol. 2018 Aug 1;202:181–90.

121. Camacho C, Coulouris G, Avagyan V, Ma N, Papadopoulos J, Bealer K, et al. BLAST+: architecture and applications. BMC Bioinformatics. 2009 Dec 15;10(1):421.

122. Robinson JT, Thorvaldsdóttir H, Winckler W, Guttman M, Lander ES, Getz G, et al. Integrative genomics viewer. Nat Biotechnol. 2011 Jan;29(1):24–6.

123. Cho KW, Youn HY, Okuda M, Satoh H, Cevario S, O’Brien SJ, et al. Cloning and mapping of cat (Felis catus) immunoglobulin and T-cell receptor genes. Immunogenetics. 1998 Jan 1;47(3):226–33.

124. Lefranc MP. From IMGT-ONTOLOGY CLASSIFICATION Axiom to IMGT Standardized Gene and Allele Nomenclature: For Immunoglobulins (IG) and T Cell Receptors (TR): Figure 1. Cold Spring Harb Protoc. 2011 Jun;2011(6):pdb.ip84.

125. Gupta NT, Vander Heiden JA, Uduman M, Gadala-Maria D, Yaari G, Kleinstein SH. Change-O: a toolkit for analyzing large-scale B cell immunoglobulin repertoire sequencing data. Bioinformatics. 2015 Oct 15;31(20):3356–8.

