## Supplementary material for "Fatal consequences of feline coronavirus infection are associated with virus persistence and a distinct adaptive immune repertoire"

**Figure S1**

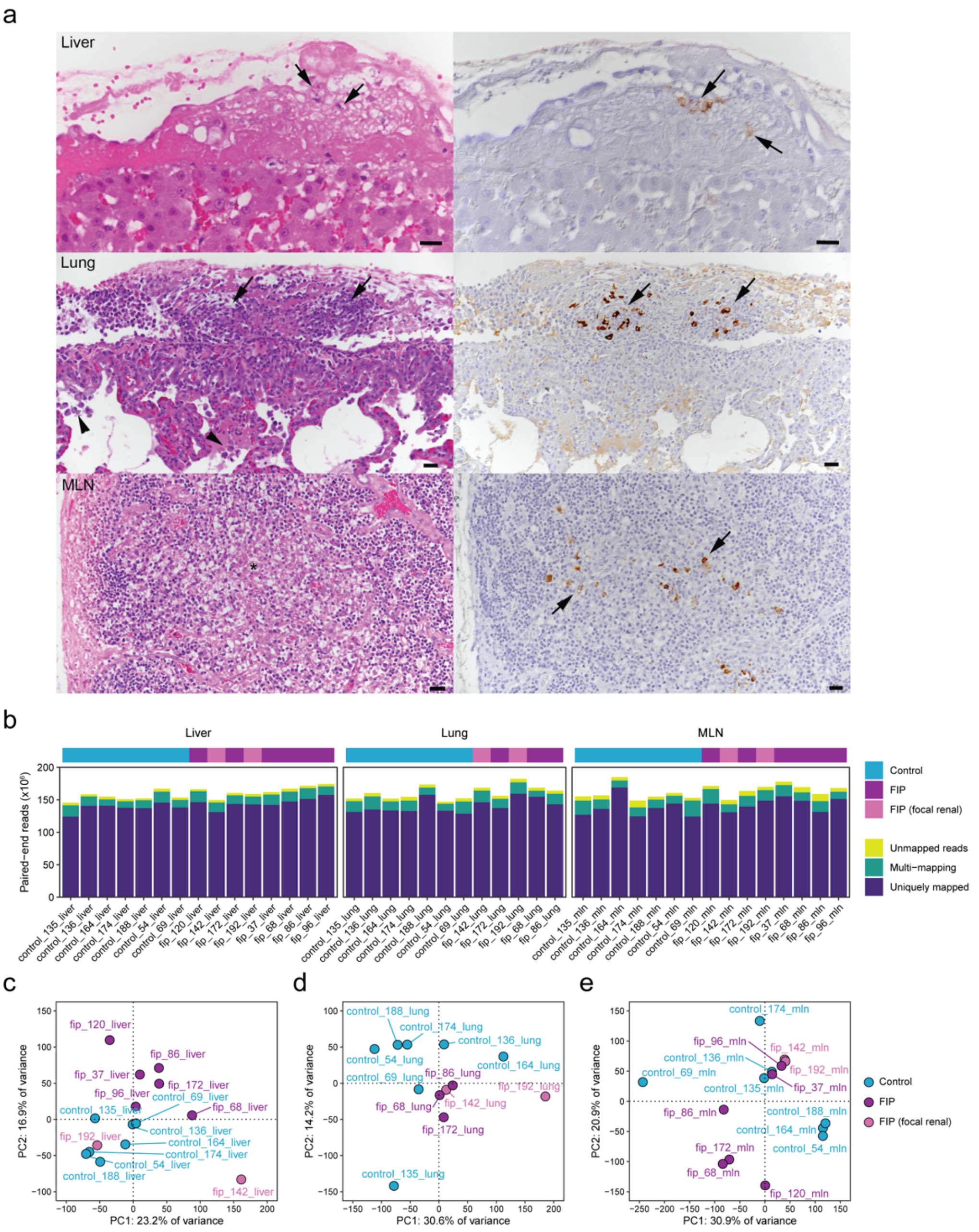


(a) Examples of histological features (left column; HE stain) and FCoV antigen (right column; IHC) within macrophages in granulomatous infiltrates. Liver (FIP cat 120) with fibrin layer on the serosal surface, with embedded macrophages (arrows) some of which express viral antigen. Bars = 25 µm. Lung (FIP cat 172) with fibrinous and granulomatous pleuritis. The granulomatous infiltrates (arrows) contain numerous strongly FCoV antigen positive macrophages. Bars = 20 µm. Mesenteric lymph node (MLN; FIP cat 120) with granulomatous infiltrate (*), containing several macrophages expressing FCoV antigen (arrows). (b) Mapping statistics for feline tissue RNA-seq reads following alignment with STAR. Bars are coloured by proportion of reads in each sample that mapped to a single unique genomic position, proportion of reads mapping to multiple loci, and proportion of reads unmapped to the felCat9.0 reference genome (GCA_000181355.4). (c-e) PCA plots based on normalised gene expression in (c) liver, (d) lung and (e) MLN samples. In each plot, each labelled point represents an individual sample, and samples are coloured by FIP status.

### Figure S2

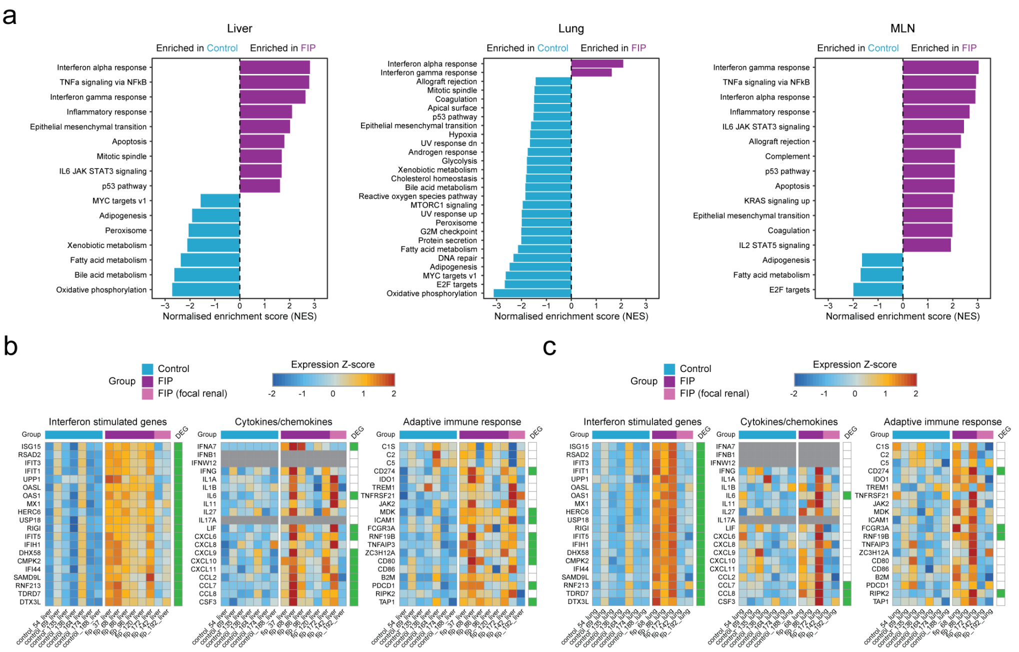


(a) Results of MSigDB feline gene set enrichment analysis (GSEA) in each tissue. All gene sets with a raw p-value and false discovery rate (FDR) < 0.05 are shown. Bars are coloured by sample group in which the gene set is enriched - FIP (purple) or control (blue). (b-c) Heatmaps showing scaled normalised expression for selected genes (rows) shown for the MLN in Fig. 2e in (b) liver and (c) lung samples (columns). Cells coloured grey indicate the gene was not detected in the tissue shown. Rows are labelled according to whether the gene was significantly differentially expressed (adjusted p-value < 0.05) in that tissue.

### Figure S3

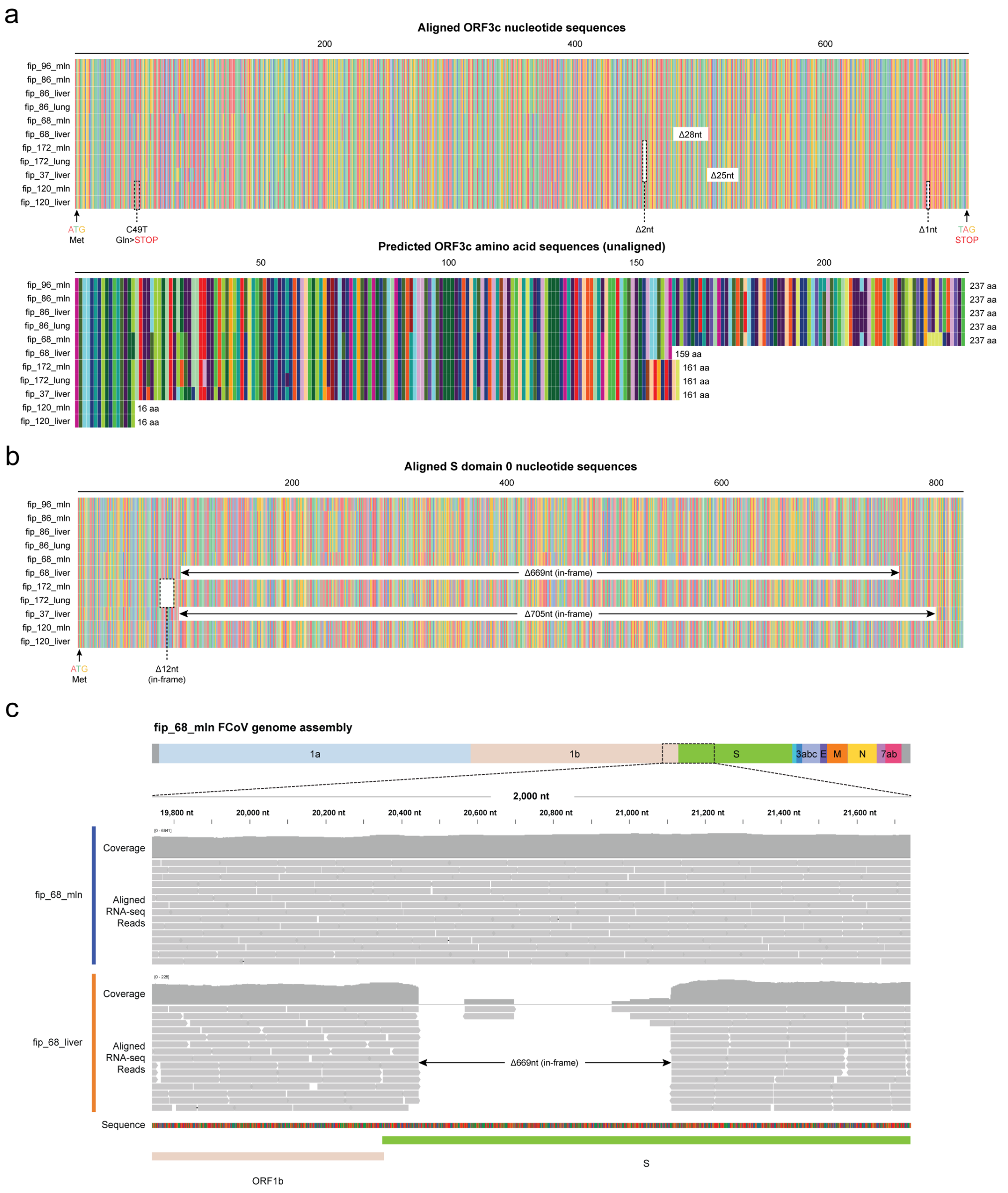


(a) Top - multiple sequence alignment of nucleotide sequences for FCoV ORF3c assembled from feline tissue RNA-seq reads. Full-length sequences are shown, from the canonical start codon to the canonical stop. Significant mutations affecting the translated product in each sample are highlighted. Bottom - unaligned amino acid sequences resulting from the FCoV ORF3c genomic sequences shown above. Each sequence is labelled with the length of the translated product from the canonical start codon to the first in-frame stop codon. (b) Top - multiple sequence alignment of nucleotide sequences for FCoV S N-terminal domain 0 assembled from feline tissue RNA-seq reads. In-frame deletions in affected samples are highlighted. (c) Integrative genomics viewer (IGV) snapshot of feline tissue RNA-seq reads from the MLN and liver of FIP cat 68 aligned to the FCoV genome assembled from the MLN sample. For each sample, the coverage is shown (log scale) above the positions of mapped reads.

### Figure S4

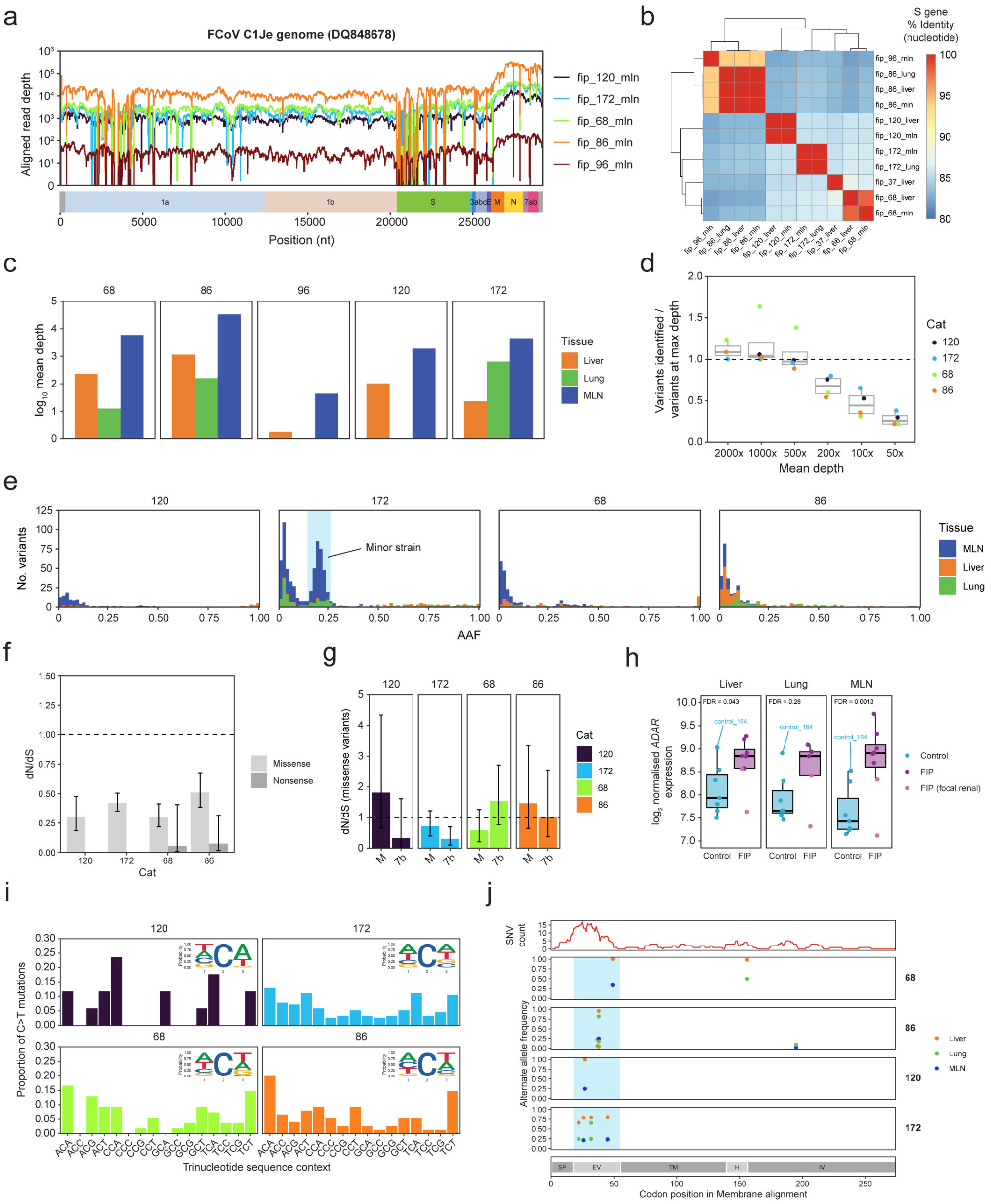


(a) Coverage of the FCoV C1Je genome (DQ848678) for feline MLN RNA-seq reads from cats for which a full-length FCoV genome was assembled using coronaSPAdes. Reads were mapped using bwa-mem, and depth at each position was determined using samtools. (b) Heatmap showing pairwise nucleotide identity (%) for S gene sequences in all full-length FCoV genomes assembled from feline tissue RNA-seq reads. (c) Log$_{10}$ mean read depth for all tissue samples from cats with a full-length FCoV genome assembled from the MLN sample. For each tissue sample, reads were mapped to the MLN FCoV genome assembly from the same cat. (d) LoFreq FCoV variants called from downsampled reads in MLN samples with high depth, relative to variants called using all aligned RNA-seq reads from the same sample. (e) Histograms showing the distribution of alternate allele frequency (AAF) of all variants called in each feline tissue sample from each cat. For FIP cat 172, a peak at an AAF of ~0.2 indicates possible co-infection with at least 2 strains of FCoV. (f) Genome-wide dN/dS estimates for missense and nonsense variants detected in each cat. Variants detected in multiple tissues were de-duplicated to retain unique variants detected in each cat. Error bars show 95% confidence intervals. For FIP cats 120 and 172, there were too few nonsense variants to accurately determine dN/dS estimates. (g) dN/dS estimates for missense variants detected in M and ORF7b genes in the FCoV genomes assembled from the MLN of each cat. Error bars show 95% confidence intervals. (h) $Log_2$ normalised expression (TMM-normalised counts per million mapped reads) of *ADAR* in each tissue determined by RNA-seq. For each tissue, the adjusted p-value/false discovery rate (FDR) derived from the edgeR differential gene expression analysis is shown. (i) Trinucleotide sequence context of C>T mutations detected in all tissue samples for each cat. Variants detected in multiple tissues were de-duplicated to retain unique variants detected in each cat. For each cat, the plot includes sequence logos showing the probability of nucleotide sequence at positions either side of the mutated residue. (j) Focused view of missense SNVs detected in multiple tissues, or at > 0.5 AAF in the liver or lung, in the M gene. The top track (red line) shows a sliding window analysis (window size = 10 residues) of the density of all SNVs detected in all samples, with adjusted codon positions from a multiple sequence alignment of predicted membrane protein sequences from the MLN FCoV assemblies of the cats shown. Each point represents an individual SNV, and points are coloured by tissue in which the SNV was detected. A variable region common to all cats, with high SNV density, is highlighted in light blue. The bottom track shows the domain structure of FCoV membrane protein, with residue numbers derived from alignment with the sequence of SARS-CoV-2 membrane protein (PDB 7VGR/7VGS). SP – signal peptide, EV – extravirion, TM – transmembrane, H – hinge, IV – intravirion.

### Figure S5

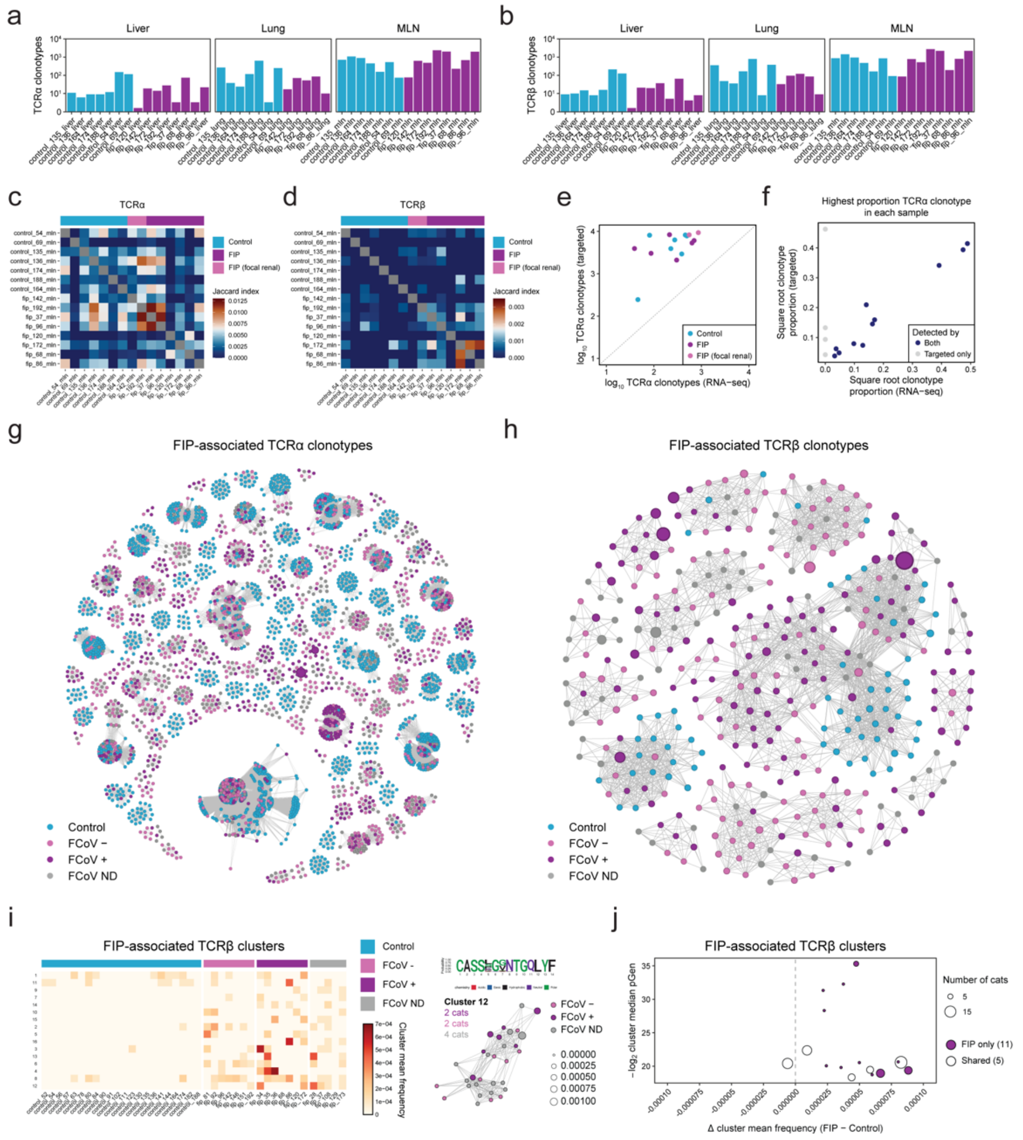


(a-b) Count of TCR clonotypes ($\log_{10}$) assembled from each RNA-seq sample for the (a) α chain and (b) β chain using MiXCR. (c-d) Jaccard index for pairwise overlap of TCR clonotypes assembled from feline tissue RNA-seq samples for the (c) α chain and (d) β chain. (e) Count of TCRα clonotypes ($\log_{10}$) assembled from either RNA-seq or targeted TCR sequencing for the MLN samples included in the RNA-seq experiment. Each point represents an individual sample, and points are coloured by FIP status. (f) Proportion of reads (square root transformed) assigned to the top TCRα clonotype (by proportion) identified by targeted TCR sequencing using either RNA-seq or targeted sequencing for each sample included in the RNA-seq experiment. Each point represents an individual sample, and samples for which the top targeted sequencing TCRα clonotype was detecting in both RNA-seq and targeted sequencing are shown in blue, and samples for which the top clonotype was detected only by targeted sequencing are shown in grey. (g-h) Networks of FIP-associated TCR clonotypes for the (g) α chain and (h) β chain detected by targeted TCR sequencing. Nodes represent unique TCR clonotypes, and edges connect clonotypes with a Hamming distance ≤ 1 in the CDR3 amino acid sequence. Nodes are coloured by FCoV status of the sample from which the clonotype was assembled, and node size is proportional to the proportion of the sample's repertoire occupied by the clonotype. (i) Left - heatmap of FIP-associated TCRβ clusters, showing the mean frequency of the cluster in each sample. Right - network plot and CDR3 sequence logo for cluster 12, with the greatest difference in cluster mean frequency between FIP and control samples. (j) Plot showing the relationship between difference in cluster mean frequency between FIP and control samples and the median cluster generation probability (pGen). Each point represents an individual TCRβ cluster, and the size of each point is proportional to the number of cats with clonotypes included in the cluster. Filled points are exclusive to FIP (purple) and unfilled points represent clusters comprised of clonotypes from both FIP and control cats.

### Figure S6

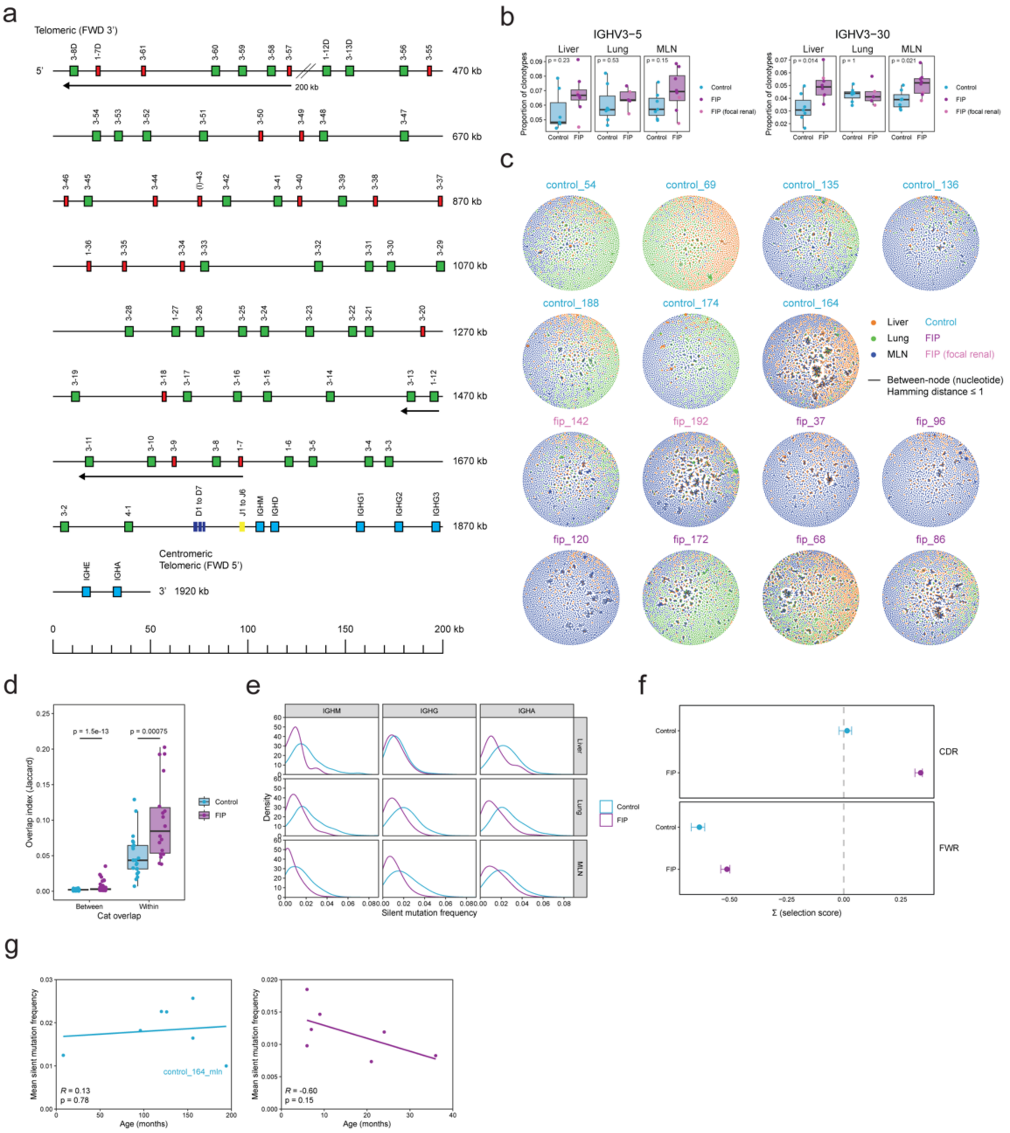


(a) IMGT-style map of the feline IGH locus, determined based on prediction of IGH VDJ and constant genes in fca126 (see Methods). (b) Boxplots showing selected IGHV gene usage in FIP and control samples. P-values were determined using the Wilcoxon test. (c) Representative networks of BCR (IGH) clonotypes assembled for tissue samples from each cat included in the RNA-seq experiment (see Methods). Each node represents a unique IGH clonotype, and edges connect nodes with a Hamming distance ≤ 1 in the CDR3 nucleotide sequence. Nodes are coloured by tissue from which the clonotype was assembled, and node size is proportional to the proportion of the sample's repertoire occupied by the clonotype. (d) Jaccard index for overlap of IGH clusters between cats with the same FIP status, and within individual cats (i.e. between tissue samples from the same cat). P-values were determined using the Wilcoxon test. (e) Density plots of the silent (non-synonymous) mutation frequency for IGH clonotypes containing the IGHM, IGHA and IGHG constant genes, across all samples from each tissue. (f) Selection strength based on the BASELINe probability density functions for IGH clonotypes using IGHG in all FIP and control samples in complementarity determining regions (CDR) and framework regions (FWR). Points represent the estimates for Σ, and error bars show the 95% confidence intervals. (g) Relationship between cat age (in months) and mean silent mutation frequency in IGH clonotypes using IGHG assembled from the MLN sample of cats included in the RNA-seq experiment. Age at the time of euthanasia was not available for FIP cat 37, so data from this cat is not shown.
